# Calcium: Modulator of Post-transcriptional and post-translational process in mESCs

**DOI:** 10.1101/2025.10.16.681898

**Authors:** Shikha Sharma, Dasaradhi Palakodeti, Tina Mukherjee

## Abstract

Calcium ions (Ca^2+^) are ubiquitous intracellular and extracellular messengers that regulate several cellular activities. Previous findings have reported active Ca^2+^ receptors in mouse embryonic stem cells (mESCs), but the fundamental requirement for Ca^2+^ signalling remains unclear. We have noted high Ca^2+^ levels in undifferentiated mESCs and observed that its depletion induces G2/M cell cycle arrest, spontaneous differentiation of mESCs towards mesoderm-like lineage, and changes in mitochondrial morphology and number. Further, our data show that Ca2+ regulates the stability of Oct4 and Nanog at the post-translational level through a pCamkIIα-dependent mechanism independent of the Jak-Stat3, transcriptional and translational control. Our data also indicate the role of Ca^2+^ at the post-transcriptional level in regulating p-bodies and stress granule markers (Dcp1a, Xrn1, Tudor, and Edc4), as well as splicing-dependent and 3’UTR-dependent NMD activity. Together, this study illustrates the broad role of Ca^2+^ in modulating key processes in mESCs.

**Highlights:**

- Ca^2+^ is present at elevated levels in mESCs and is important for pluripotency
- Ca^2+^ depletion accelerates G2/M cell cycle arrest and mesoderm-like traits
- Ca^2+^ regulates pluripotency and p-bodies markers at the post-translational level
- Ca^2+^ regulates Oct4, Nanog, Dcp1a through pCamkIIα dependent mechanism

## Introduction

Embryonic stem cells (ESCs) are derived from the inner cell mass (ICM) of embryos and are characterised by pluripotency, self-renewal, and limitless proliferation (Ye et al., 2024). Several studies have shown that the ESCs’ core pluripotency network is regulated by diverse molecular mechanisms, extracellular signalling pathways, post-transcriptional regulation, epigenetic mechanisms, and chromatin remodelling (Yeo and Ng, 2013, Kobayashi and Kikyo, 2015). The ability of ESCs to differentiate into cells from the three embryonic germ layers makes them ideal candidates for basic research on pluripotency, genomic engineering, cell identity transitions, disease modelling, cell therapy, and regenerative medicine (Kobayashi and Kikyo, 2015, Czechanski et al., 2014). Further understanding the regulation of ESCs’ pluripotency network will provide more insight into the nature of pluripotent stem cells, cell fate decision, and lineage specification.

Calcium (Ca^2+^) is a fundamental regulator of several processes such as cell cycle, cell proliferation, neuronal plasticity, gene regulation, and apoptosis (Clapham, 2007, Berridge et al., 2003). Despite the broad role of Ca^2+^ signalling in different cell types, there are only a few studies that have reported the presence of Ca^2+^ signalling in pluripotent stem cells. In 2004, Yanagida et al. documented the activation of various Ca^2+^ receptors upon stimulation in mESCs, indicating the presence of calcium-regulating pathways (Yanagida et al., 2004). Todorova et al. (2009), observed the activation of Ca^2+^ signalling by lysophosphatidic acid (LPA) in mESCs. In 2010, Schwirtlich et al. reported that GABA-mediated Ca^2+^ signalling is important for the proliferation and differentiation of mESCs. Recently, it was demonstrated that high intracellular calcium is paramount for regulating naïve pluripotent state and self-renewal of mESCs, suggesting the importance of calcium in controlling the pluripotency of mESCs (Zhu and Zhang, 2019, MacDougall et al., 2019). However, the role of Ca^2+^signalling in the regulation of pluripotency remains unclear. In this study, we have reported the role of Ca^2+^ signalling in regulating proliferation, cell cycle, p-bodies dynamics and core pluripotency markers of mESCs. Our data indicates that high Ca^2+^ is important for the maintenance of pluripotency in mESCs at the post-transcriptional and post-translational levels.

## Results

### Pluripotent stem cells maintain an elevated level of cytosolic calcium

Exogenous addition of leukaemia inhibitory factor (LIF) is crucial for the maintenance of mESCs pluripotency (Ohtsuka et al., 2015, Godwin et al., 2017). Using Fluo-4 staining, we observed a high level of Ca^2+^ in undifferentiated mESCs cultured in control media containing LIF (C+L) compared to spontaneously differentiated mESCs cultured in the absence of LIF (C-L) (Fig. 1A and 1D). It is not yet clear why undifferentiated mESCs maintain a high level of Ca^2+^ within their cytoplasm. Therefore, we became interested in understanding the role of Ca^2+^ in mESCs. We manipulated the level of Ca^2+^ in the presence of LIF using BAPTA-AM (BAPTA) and thapsigargin. BAPTA is a highly selective, cell-permeable Ca^2+^ chelator compared with other Ca^2+^ chelating agents, including EGTA, EDTA, and Fura-2, and is widely used in hundreds of studies to explore Ca2+ signalling (Collatz et al., 1997; Sneyers et al., 2024). Thapsigargin is a selective SERCA pump inhibitor that does not affect other ion-motive ATPases, such as plasma membrane receptors, and is known to increase intracellular Ca^2+^ in various cell types, including ESCs (Lytton et al., 1991; Huang et al., 2017; Thomas and Hanley, 1994; Taylor and Broad, 1998). We checked the viability of mESCs using the Trypan Blue exclusion test (Strober 2015). We observed that all cells were viable across different concentrations of BAPTA (1μM, 2μM, and 5μM) and thapsigargin (250 pM, 500 pM, and 1nM) (data not shown), indicating no apoptosis. Using Fluo-4 staining, we observed that BAPTA 2μM (B2μM) supplementation for 72 hours reduced Ca^2+^ levels in mESCs to levels comparable to the no-LIF condition (C-L), whereas thapsigargin 1nM (T1nM) treatment increased Ca2+ levels (Fig. 1A and 1D). We also noted that Ca^2+^ was downregulated in the C-L condition, and T1nM supplementation in the C-L condition slightly increased Ca^2+^ levels but did not restore them to the control condition (C+L) (Fig. 1A and 1D). Similarly, using GCaMP6f, we observed increased Ca2+ levels after T1nM treatment and decreased levels after B2μM treatment and in the C-L condition (Supp Fig. 1A and 1B). We also detected heterogeneity in the level of Ca^2+^ in cells within mESCs colonies (Fig. 1A). In addition, we noted that B2μM also reduced the Ca^2+^ level in the endoplasmic reticulum (ER) (Fig. 1B and 1E) and peri-lysosomes (Fig. 1C) of mESCs. Conversely, T1nM increased the overall Ca^2+^ level in the peri-lysosomes (Fig. 1C) but not in the ER (Fig. 1B and 1E). Indeed, the peri-lysosomal region has been implicated in calcium signalling and autophagy (Nguyen et al., 2024; Saffi et al., 2024).

**Figure 1:**
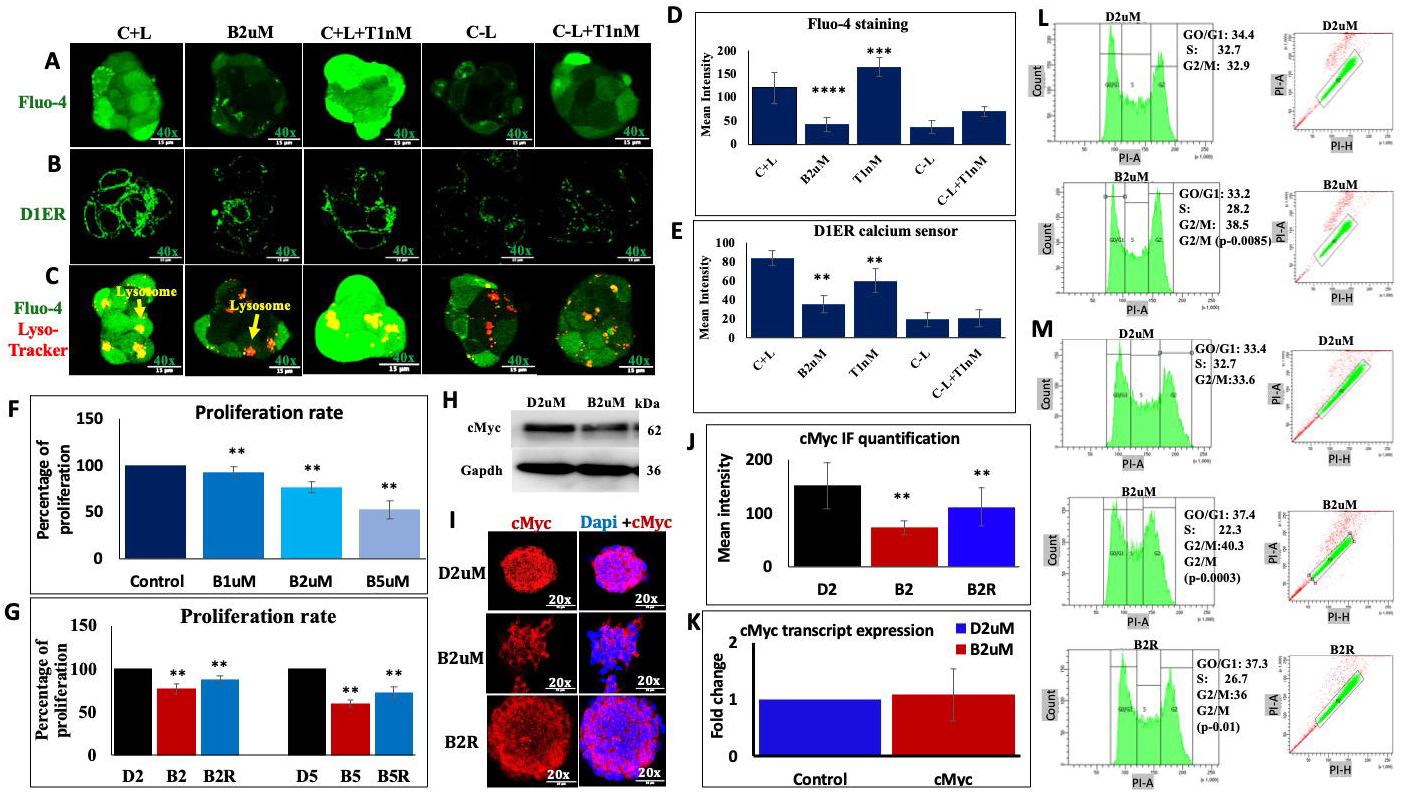
Calcium level detection in mESCs: **A)** Fluo-4 staining was done to measure intracellular Ca^2+^ level using excitation 488nm laser, B) D1ER reporter expression for the detection of Ca^2+^level in endoplasmic reticulum (ER) (green) using excitation 488nm laser, C) co-staining of Fluo-4 (green) and lyso Tracker deep red (red) for the detection of peri-lysosomal Ca^2+^ level using 488 and 633nm laser, in mESCs under different conditions (C+L, B2μM, T1nM, C-L, C-L+T1nM) (n=3-5) obtained at 40x magnification with scale bar 15μm shown by representative images. **D and E)** Bar plot represents the mean intensity of Fluo-4 staining with B2μM, C+L+T1nM compared to control, E) mean intensity of D1ER reporter in mESCs with B2μM and C+L+T1nM compared to control. The mean intensity was calculated from mESCs colonies containing approximately 11-23 cells from the confocal images (n=12) from different region of the above experiments. **F)** Bar graph showing the percentage of the proliferation of mESCs treated with B1μM, B2μM, and B5μM for 72 hrs compared to control using the Trypan Blue method (n=4). **G)** Bar graph showing percentage of the proliferation of mESCs treated with B2μM and B5μM for 5 days and B2R condition compared to control D2 and D5, respectively (n=4). **H)** Western blot detection of cMyc in the presence and absence of B2μM for 72 hours (n=3) shown by representative images. Protein estimation was done using the Bradford method, and an equal amount of total protein from the cell lysate was loaded into the SDS-PAGE gel for all the samples. **I)** Immunofluorescence (IF) detection of cMyc expression in mESCs in the presence of B2μM for 5 days and B2R condition (n=3) at 20x magnification with scale bar 35μm shown by representative images. IF experiments were done by thorough scanning the slide of different areas. **J)** The bar plot represents the mean intensity of cMyc expression in mESCs colonies containing 20-70 cells, calculated from the confocal images (n=12) from different regions of the above experiments. **K)** QPCR of cMyc expression in mESCs treated with B2μM for 72 hours (n=3). **L)** Cell cycle analysis of control (D2) and B2μM treated mESCs for 72 hours (n=4). A significant p-value (0.0085) was detected for B2/D2 (G2/M). The left panel is a linear DNA count histogram. The right panel shows the pulse geometry gating for doublet discrimination. Gating was done on the linear diagonal cluster of single cells to exclude cell aggregates and the apoptotic population. **M)** Cell cycle analysis of D2, B2μM treated mESCs for 5 days and B2R condition (n=5). Significant p-value (0.0003 and 0.01) was detected for B2/D2 and B2R/B2 (G2/M). The left panel is a linear DNA count histogram. The right panel shows the pulse geometry gating for doublet discrimination. Gating was done on the linear diagonal cluster of single cells to exclude cell aggregates and the apoptotic population. *C+L= Control media containing Lif; B1*μ*M= C+L + BAPTA 1*μ*M; B2*μ*M = C+L + BAPTA 2*μ*M; T1nM= C+L + thapsigargin 1nM; C-L= control media without Lif; C-L+T1nM = C-L + thapsigargin 1nM; D2= C+L+ DMSO equivalent to B2*μ*M condition; B5*μ*M = C+L+ BAPTA 5*μ*M; D5= C+L+ DMSO equivalent to B5*μ*M condition, B2R= B2 media for 72 hour + D2 media for 48 hour, n = biological replicates, BAPTA and thapsigargin were used as calcium-modulating reagents*. *The mean intensity was calculated from the confocal images from different regions for all experiments using ImageJ software.* An unpaired two-sample t-test was used to calculate the p-value using the MedCalc online statistical software calculator (MedCalc Software, 2024) with the mean and standard deviation. ***p-value representation with asterisk <0.05 (*), <0.01 (**), <0.001 (***), <0.0001 (****)*.**

We also assessed proliferation using the Trypan Blue exclusion test (Funk et al., 2021; Wiepz et al., 2006). We observed a decrease in the proliferation of mESCs with increasing concentrations of BAPTA (B1μM, B2μM, and B5μM) after 72 hrs (Fig. 1F). We noted that mESCs colonies exhibited fibroblast morphology upon differentiation and observed increased fibroblast morphology with increasing concentrations of BAPTA (B1μM and B2μM) (Supplementary file S1 (S1)_Supplementary Fig. S3C and S3D). We observed that undifferentiated mESCs formed characteristic dome-shaped colonies with tightly packed cells; B2μM-treated mESCs exhibited loose dome-shaped colonies that spread into a monolayer of fibroblast-like cells; and the C-L condition showed an extensive monolayer of fibroblast-like cells, indicating less differentiation in BAPTA-treated samples than in C-L (S1_ Supplementary Fig. S3C and S3D). We further replaced B2μM media after 72 hours with control ESC media (B2R) for 48 hours. We observed an increase in the proliferation of mESCs, but not to the extent of the control condition, indicating that BAPTA treatment is partially reversible (Fig. 1G). However, we noted that mESCs that underwent complete morphological change (fibroblast appearance) could not proliferate further and retained their phenotype. Myc transcription factor plays an important role in ESC self-renewal and pluripotency (Fagnocchi et al., 2016). Several studies have shown that Myc is important for maintaining unlimited mESCs proliferation, and its depletion induces dormancy and proliferation arrest (Scognamiglio et al., 2016; Fagnocchi et al., 2016; Ruan et al., 2017). c-Myc is regarded as the key player to maintain undifferentiated state and ESCs self-renewal (Hishida et al., 2011). We next assessed cMyc levels in mESCs. We observed that cMyc expression decreases upon B2μM treatment after 72 hours by western blot and immunofluorescence (IF) (Fig. 1H, 1I, 1J), independent of transcriptional control (Fig. 1K), which indicated that Ca^2+^ might be regulating proliferation/stemness in mESCs by controlling cMyc expression. We also observed increased cMyc expression in B2R compared with B2μM at 5 days (Fig. 1I and 1J). However, we were unable to study the role of N-myc upon BAPTA treatment; thus, further studies are needed to explore the role of N-myc and other proteins crucial for mESCs self-renewal upon calcium manipulation. In addition, we analysed the cell cycle profile of C+L, B2μM, and B2R mESCs at 72 hrs (Fig. 1L) and 5 days (Fig. 1 M) and observed that B2μM treatment caused cell cycle arrest at G2/M, which was reversible with B2R treatment (Fig. 1M). Earlier reports in mesenchymal stem cells (MSCs) and epidermal stem cells suggest that high calcium is crucial for stem cell proliferation and cell cycle progression, and that the G2 phase of the cell cycle is important for determining the Ca2+ signalling pattern (Moore et al., 2024; Vallet et al., 2025). Further, we also noted that B2μM and C-L resulted in fluctuations in the expression of calcium receptors of the plasma membrane, ER and mitochondria, including *Pmca1, Pmca2, Pmca3 and Pmca4; Serca1, Serca2 and Serca3; Itpr1, Itpr2, and Itpr3; Stim1 and Stim2; Orai1 and Orai3; Ryr1 and Ryr2 and Mcu* (S1_Supplementary Fig. S2C, 2D, 2E, 2F, 2G, 2H), which signifies that BAPTA treatment might be affecting the level of Ca^2+^ within different organelles.

**Figure 2:**
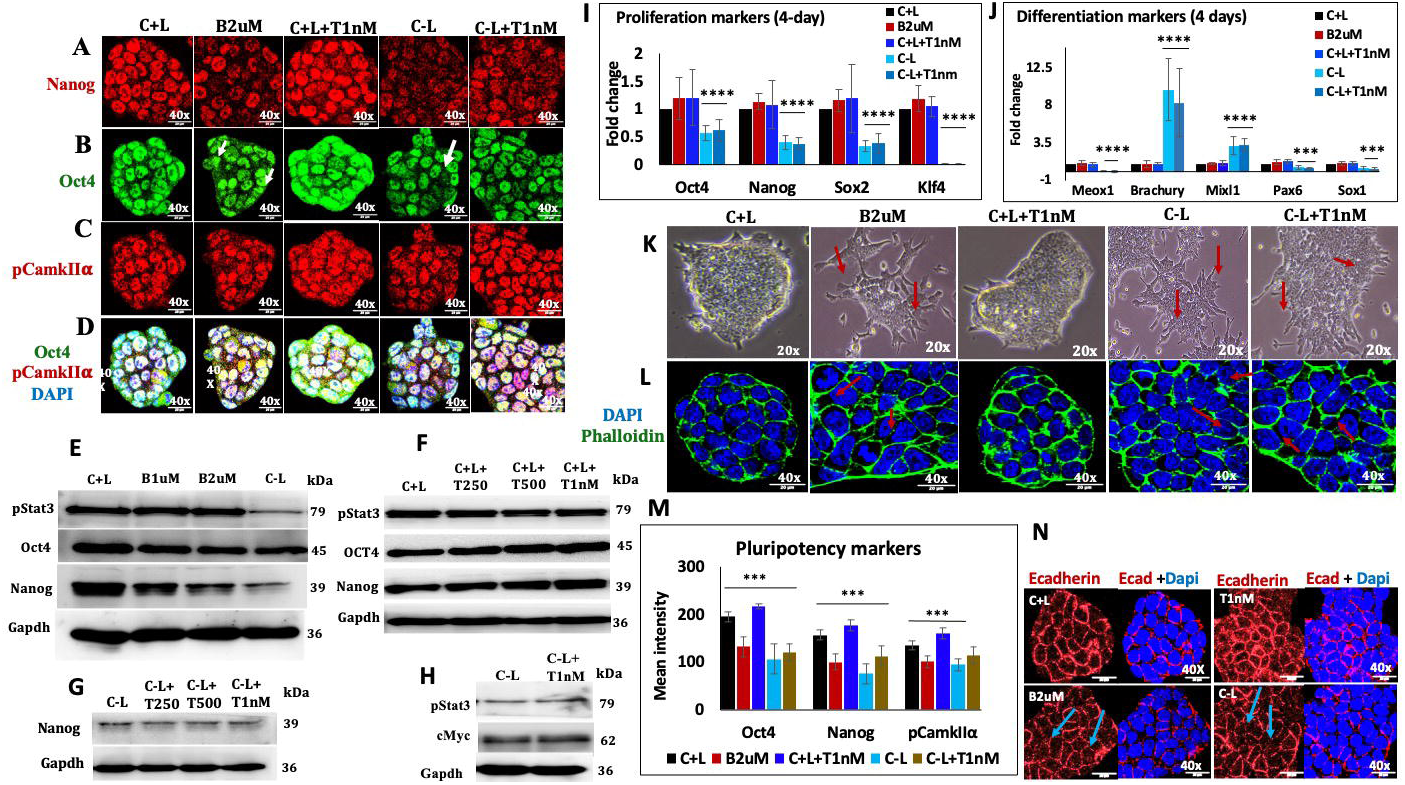
Role of calcium in pluripotency: Immunofluorescence (IF) detection of **A)** Nanog (red), **B)** Oct4 (green), **C)** pCamkIIα (red), **D)** Oct4, pCamkIIα and DAPI co-staining in mESCs cultured under different conditions (C+L, B2μM, T1nM, C-L, C-L+T1nM) (n=3) at 40x magnification with a 20 μm scale bar shown by representative images. IF experiments were done by thorough scanning the slide of different areas. **E)** Western blot shows the expression of Oct4, Nanog, pStat3 and Gapdh in mESCs cultured under different conditions (C+L, B1μM, B2μM, C-L) (n=3) shown by representative images. Protein estimation was done using the Bradford method, and an equal amount of protein from the cell lysate was loaded into the SDS-PAGE gel for all the samples. **F)** Western blot shows the expression of Oct4, Nanog, pStat3 and Gapdh in mESCs cultured under different conditions (C+L+T250 pM, C+L+T500 pM, C+L+T1nM) (n=3) shown by representative images. Protein estimation was done using the Bradford method, and equal amounts of total protein from the cell lysate were loaded into the SDS-PAGE gel for all the samples. **G)** Western blot showed expression of Nanog in mESCs cultured under different conditions (C-L, C-L+T250pM, C-L+T500pM, C-L+T1nM) (n=3) shown by representative images. Protein estimation was done using the Bradford method, and equal amounts of total protein from the cell lysate were loaded into the SDS-PAGE gel for all the samples. **H)** Western blot showed expression of pStat3 and cMyc in mESCs cultured under different conditions (C-L, C-L+T 1nM) (n=3) shown by representative images. Protein estimation was done using the Bradford method, and equal amounts of total protein from the cell lysate were loaded into the SDS-PAGE gel for all the samples. **I, J)** QPCR of pluripotency markers and early differentiation markers under different conditions (C+L, B2μM, T1nM, C-L, C-L+T nM) (n=5). **K)** Brightfield images of the mESCs cultured under different conditions (C+L, B2μM, T1nM, C-L, C-L+T1nM) (n=5) at 20x magnification. **L)** Immunofluorescence (IF) detection of Phalloidin in mESCs cultured under different conditions (C+L, B2μM, T1nM, C-L, C-L+T1nM). (n=3) at 40x magnification with 20μm scale bar shown by representative images. **M)** The bar graph represents the mean intensity of Oct4, Nanog and pCamkIIα expression calculated for mESCs colonies containing approximately 20-45 cells from the confocal images (n=15) from the different regions of the above experiments (A, B, C). **N)** Immunofluorescence (IF) detection of E-cadherin staining in mESCs cultured under different conditions (C+L, B2μM, T1nM, C-L) (n=3) at 40x magnification with a 20 μm scale bar shown by representative images. IF experiments were done by thorough scanning the slide of different areas. ***Note**: C+L = Control media containing Lif; B1*μ*M = C+L + BAPTA 1* μ*M; B2*μ*M = C+L + BAPTA 2* μ*M; T1 nM = C+L + thapsigargin 1 nM; C-L = control media without Lif; C-L+T1 nM = C-L + thapsigargin 1 nM; T250 pM, T500 pM, and T nM = C+L + thapsigargin (250 pM, 500 pM, and 1 nM), n= biological replicates. BAPTA and thapsigargin were used as calcium-modulating reagents*. *The mean intensity was calculated from the confocal images from different regions for all experiments using ImageJ software.* An unpaired two-sample t-test was used to calculate the p-value using the MedCalc online statistical software calculator (MedCalc Software, 2024) with the mean and standard deviation. ***p-value representation with asterisk <0.05 (*), <0.01 (**), <0.001 (***), <0.0001 (****)*.**

### Calcium homeostasis is critical for pluripotency

To explore the role of Ca^2+^ in maintaining pluripotency in mESCs, we assessed pluripotency marker expression after BAPTA and thapsigargin treatment. Western blot and immunofluorescence (IF) data confirmed the increase/decrease in the protein level of core pluripotency markers including Oct4 and Nanog upon low/high Ca2+ condition using BAPTA (B1μM and B2μM) and thapsigargin 250 pM, 500 pM and 1nM (T250 pM, T500 pM, T1nM) in the presence of LIF, suggesting that Ca^2+^ has role in the regulation of pluripotency markers (Fig. 2 A, B, E, F and M). Moreover, we noted the heterogeneity in the expression of Oct4 and Nanog in cells within mESCs colonies (Fig. 2A and 2B), which also corroborates the heterogeneous level of Ca^2+^ in cells within mESCs (Fig. 1A). In addition, we have not observed any significant difference in the expression of Oct4, Nanog, and cMyc in the absence of LIF and in the presence of thapsigargin (Fig. 2 A, B, G, H, and M). CHIR 99021 (CH) and PD0325901 (PD) also known as 2i inhibitors, inhibit glycogen synthase kinase 3 beta (GSK3β) and MAPK/ERK signalling pathways, implicated in maintaining the ground state of mESCs [30] (Sim et al., 2017). Furthermore, we treated mESCs with 2i inhibitors in the presence of LIF and B1μM and observed decreased expression of Oct4 and Nanog proteins (S1_Supplementary Fig. S3A), indicating that Ca^2+^regulates pluripotency marker expression in ground-state mESCs. In addition, we observed that some of the mESCs colonies exhibited fibroblast morphology upon B1μM treatment in the presence of a 2i inhibitor (S1_Supplementary Fig. 3B).

**Figure 3:**
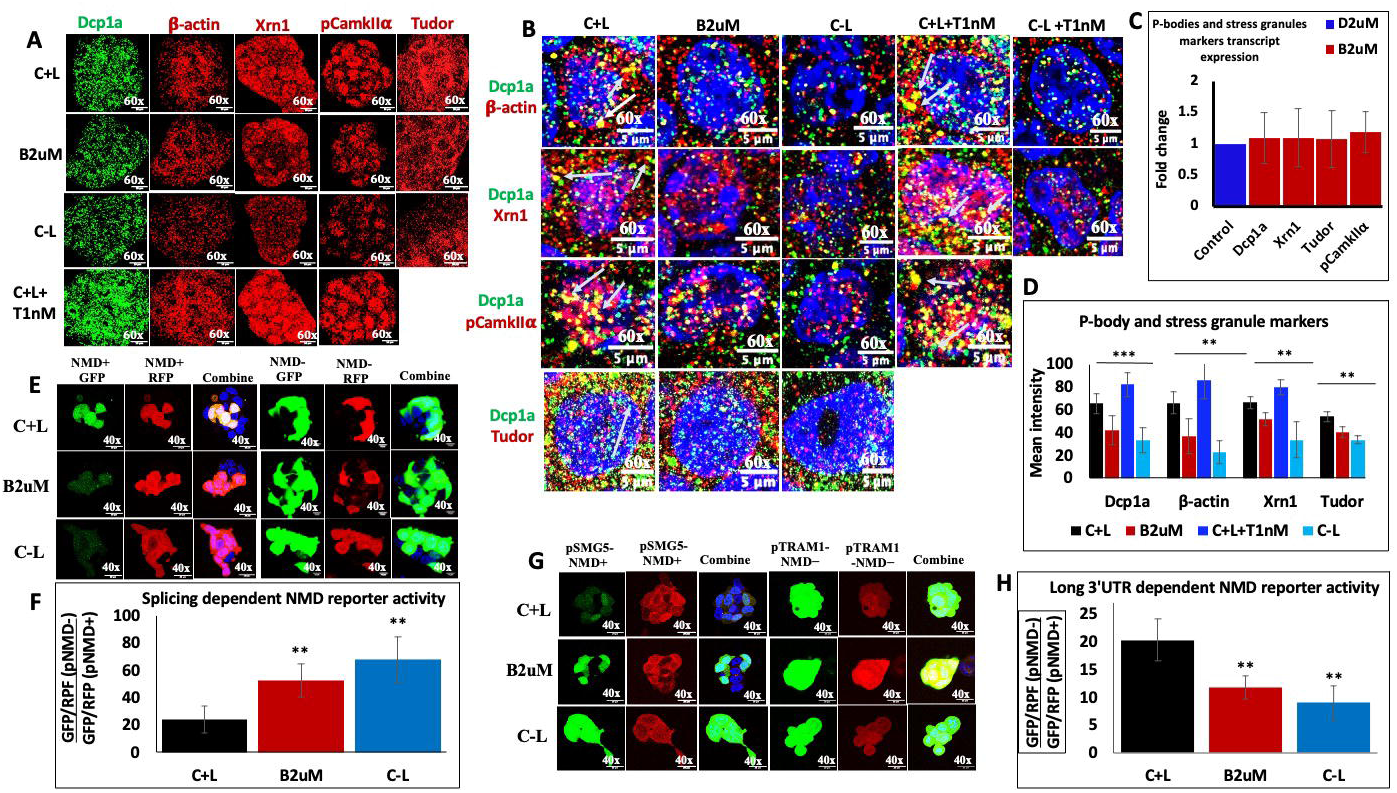
Role of calcium at the post-transcriptional level: **A)** Immunofluorescence (IF) detection of Dcp1a, β-actin, Xrn1a, pCamkIIα, Tudor in mESCs cultured under different conditions (C+L, B2μM, T1nM, C-L) (n=3-5) using confocal microscopy at 60x magnification with 10μm scale bar shown by representative images. IF experiments were done by thorough scanning the slide of different areas. **B)** Co-localization expression profile of Dcp1a and β-actin; Dcp1a and Xrn1; Dcp1a and pCamkIIα; Dcp1a and Tudor in mESCs cultured under different conditions (C+L, B2μM, T1nM, C-L, C-L+T1 nM) (n=3-5). To observe granule size and co-localization of p-bodies and stress granule markers, the images obtained using a confocal microscope at 60x magnification were cropped to one nucleus size and normalized using a 5μm scale bar at the bottom. **C)** QPCR of p-bodies and stress granule markers (Dcp1a, Xrn1, Tudor and pCamkIIα) in the mESCs cultured in the presence of B2μM for 72 hours (n=3). **D)** The bar graph represents the mean intensity of Dcp1a, Xrn1, pCamkIIα and Tudor expression in mESCs colonies containing approximately 11-20 cells, calculated from confocal images (n=15) from different regions from the above experiments (A). **E)** Immunofluorescence (IF) detection of splicing-dependent NMD by using NMD+ GFP, NMD+RFP, NMD-GFP, NMD-RFP reporter in mESCs cultured under different conditions (C+L, B2μM, C-L) (n=3) at 40x magnification with 20μm scale bar shown by representative images. **F)** Splicing-dependent NMD activity was calculated for mESCs colonies containing 5-15 cells from the confocal images (n=11) from different regions of the above experiments as described by Pereverzev et al. 2015. **G)** Immunofluorescence (IF) detection of long 3’ UTR-dependent NMD by using SMG+ GFP, SMG+RFP, pTRAM-GFP, pTRAM-RFP reporter in mESCs cultured under different conditions (C+L, B2μM, C-L) (n=3) at 40x magnification with 20μm scale bar shown by representative images. **H)** 3’ UTR-dependent NMD activity was calculated for mESCs colonies containing 5-15 cells from the confocal images (n=11) from different regions of the above experiments as described by Pereverzev et al. 2015. ***Note:*** *C+L= Control media containing Lif; B2*μ*M = C+ L + Bapta 2*μ*M; T1 nM= C+L + thapsigargin 1nM; C-L= control media without Lif; C-L+T1nM = C-L + thapsigargin 1nM, n=biological replicates, BAPTA and thapsigargin were used as calcium-modulating reagents*. *The mean intensity was calculated from the confocal images obtained from different regions for all experiments using ImageJ software.* An unpaired two-sample t-test was used to calculate p-value using the MedCalc online statistical software calculator (MedCalc Software, 2024) with the mean and standard deviation. ***p-value representation with asterisk*** *<0.05 (*), <0.01 (**), <0.001 (***), <0.0001 (****)*.

Phosphorylated CamkIIα (pCamkIIα) is an eminent downstream target of Ca^2+^ signalling (Clapham 2007, Chi et al., 2016). We checked the expression of pCamkIIα using the pCamkIIα (Thr 286) monoclonal antibody. Our data showed Ca2+-dependent simultaneous upregulation/downregulation of pCamkIIα and Oct4 in mESCs upon B2μM and T1nM treatment for 72 hours (Fig. 2B, 2C, and 2M). We further noted that pCamkIIα is present mainly in the nucleus of mESCs and observed the co-localisation of pCamkIIα with Oct4 (Fig. 2B, 2C, 2D, S1_Supplementary Fig. 4A and 4B), indicating that Ca^2+^ might have a role in the regulation of phosphorylation of pluripotency markers through a pCamkIIα-dependent mechanism. Previous reports suggest that LIF plays a major role in maintaining the pluripotency of mESCs mainly through the Jak-Stat3 pathway [12, 32] (Tang and Tian, 2013, Ohtsuka et al., 2015). We observed that the protein level of pStat3 was unchanged upon B1μM, B2μM, T250 pM, T500 pM, and T1nM treatment in the presence and absence of LIF, suggesting that Ca^2+^ might not be playing a role through the Jak-Stat3 pathway for the maintenance of pluripotency of mESCs (Fig. 2 E, F, and H). Further, we checked the transcript level of *Oct4, Nanog, Sox2 and Klf4*, and early differentiation markers of mesoderm (*Mixl1, Brachyury*), endoderm (*Sox17, and MeoX1*), and ectoderm (*Sox1, and Pax6*) upon Ca^2+^ manipulation using B2μM and T1nM and observed no change in the transcript level of these markers in 48hrs (S1_Sipplementary Fig. 2A and 2B) and 4 days (Fig. 2I and 2J). Among these early differentiation markers, we found that *Sox17* was either not expressed or showed a CT value of 35 under all conditions. Taken together, our data indicate that Ca^2+^ may play a role in regulating the protein of pluripotency and differentiation markers at the post-transcriptional (sequestering transcript in mRNP granules), translational, and post-translational levels.

**Figure 4:**
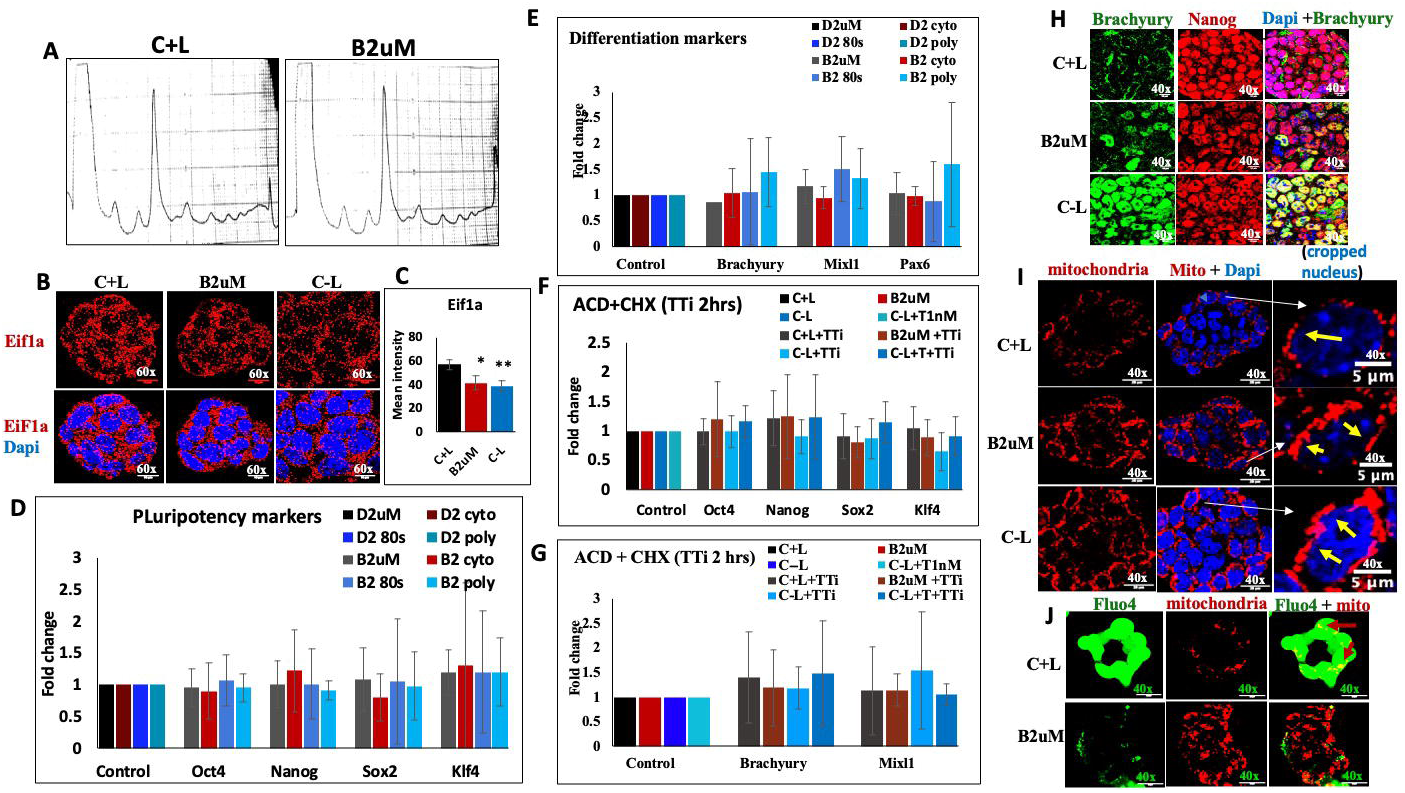
Role of calcium at translational level: A) Polysome profile of C+L and B2μM treated mESCs. (n=5). B) Immunofluorescence (IF) detection of Eif1a expression in mESCs cultured under different conditions (C+L, B2μM, C-L) using confocal microscopy at 60x magnification with 10μm scale bar. (n=3) shown by representative images. IF experiments were done by thorough scanning the slide of different areas. C) The bar graph represents the mean intensity of Eif1a expression in mESCs colonies containing approximately 15-20 cells, calculated from the confocal images (n=12) from different regions of the above experiments. D, E) QPCR of core proliferation (*Oct4, Nanog, Sox2, Klf4*) and early differentiation (*Brachyury, Mixl1, Pax6*) markers in the cytoplasmic, 80s, polysome fraction of polysome profiling of control (D2) and B2μM treated mESCs. (n=4). F, G) QPCR of core proliferation markers (*Oct4, Nanog, Sox2, Klf4*) and early differentiation markers *(Brachyury and Mixl1)* of mESCs cultured under different conditions (C+L, B2μM, C-L, C-L+T 1nM) for 72 hours followed by combined cycloheximide (CHX) and actinomycin-D (ACD) treatment (TTi) for 2 hours (C+L+TTi, B2μM+TTi, C-L+TTi, C-L+T+TTi). (n=5). H) Immunofluorescence (IF) detection of Brachyury and Nanog expression in mESCs cultured under different conditions (C+L, B2μM, C-L) at 60x magnification with 10um scale bar shown by representative images. (n=3). I) Confocal detection of mitochondria using Mito Tracker-red staining in mESCs cultured under different conditions (C+L, B2μM, C-L) for 48 hours using confocal microscopy at 40x magnification with a 20 μm scale bar shown by representative images. To observe the mitochondria shape, images were cropped to one nucleus and normalized using a 5μm scale bar at the bottom. (n=4). J) Confocal detection of mitochondria and calcium level using Mito Tracker-red and Fluo-4 co-staining in mESCs cultured under condition (C+L and B2μM) (n=3) at 40x magnification with 20μm scale bar shown by representative images. ***Note:*** *C+L= Control media containing Lif; B2*μ*M = C+ L + Bapta 2*μ*M; C-L= control media without Lif; C-L+T1nM = C-L + thapsigargin 1nM; TTi= Cycloheximide (CHX) and actinomycin-D (ACD) treatment, n= biological replicates, BAPTA and thapsigargin were used as calcium-modulating reagents*. *The mean intensity was calculated from the confocal images collected from different regions for all experiments using ImageJ software. An unpaired two-sample t-test was used to calculate the p-value using MedCalc online statistical software calculator (MedCalc Software, 2024) with the mean and standard deviation. **p-value representation with asterisk** <0.05 (), <0.01 (), <0.001 (), <0.0001 (****)*.

Besides this, we observed an increase in the cytoplasm-to-nucleus ratio of mESCs by decreasing the level of Ca^2+^ using B2μM and C-L conditions, and a decrease in the cytoplasm-to-nucleus ratio upon T1nM treatment by phalloidin staining, as shown by the arrow (Fig. 2K and 2L). Park et al. also showed that high cytosolic Ca^2+^ decreases the cytoplasm/nucleus ratio in Drosophila C4da neurons (Park et al., 2020), corroborating our findings and suggesting a possible role for Ca^2+^ in cytoskeletal remodelling during cell-shape transitions. The cadherin-adhesion complex is important for bridging cell-to-cell contact and drives various morphogenetic changes (Mege and Ishiyama, 2017). Therefore, we checked the expression of the calcium-dependent adhesion protein E-cadherin and found that E-cadherin formed membrane-like puncta around cells and that its expression was downregulated in mESCs upon B2μM treatment and C-L condition, and increased upon T1nM condition shown by arrow (Fig. 2N), indicating the involvement of Ca^2+^-dependent cadherin family proteins for driving morphological changes in ESCs upon Ca^2+^ manipulation (Fig. 2N). Spencer et al. (2007) [35] also reported decreased E-cadherin expression in spontaneously differentiated mESCs, which supports our findings.

### Calcium is important for the regulation of ribonucleoprotein complexes and NMD activity

To understand whether Ca^2+^ has any role at the post-transcriptional level. We examined the expression of P-body and stress granule markers upon Ca2+ manipulation in mESCs. Our data showed that Dcp1a, Xrn1, Tudor and β-actin are present abundantly in both the cytoplasm and nucleus of undifferentiated mESCs (Fig. 3A and 3B, S1_Supplementary Fig. S5A, 5B, 5C, and 5D). Previous studies have also reported the presence of Dcp1a in the nucleus and cytoplasm of mESCs and human cells (Jia et al., 2012; Rzeczkowski et al., 2011), which was earlier recognised as predominantly cytoplasmic. Further, we noted a decrease/increase in the expression of these mRNP granules upon B2μM and T1nM treatment (Fig. 3A, B, D, S1_Supplementary Fig. S5A, 5B, 5C, 5D, 5E). We also noted the variation in the size of these granules, as some were large in size and others small (Fig. 3B). The variations in size could be due to several reasons, such as a) these markers may be a part of several complexes having different compositions, either present as a monomer or dimer; b) they may be at different stages of maturation for post-transcriptional processes to carry out their functions. Moreover, studies in yeast showed that Ca^2+^ is required for P-body formation and revealed alterations in P-body granule size (Kilchert et al., 2010; Mark et al., 2016). Further, we have observed the co-localisation of Dcp1a with β-actin, Xrn1, Tudor, and pCamkIIα, indicating that Ca^2+^ might be regulating p-body markers through a pCamkIIα-dependent mechanism (Fig. 3B). However, we were unable to check the co-localisation of other markers with pCamkIIα as both the antibodies were raised in the same species. Our data also show that Ca^2+^ regulates P-bodies and stress granule markers independent of transcriptional control (Fig. 3C). Next, we have quantified splicing-dependent nonsense-mediated mRNA decay (NMD) activity and long 3’ UTR-dependent NMD activity in B2μM and C-L conditions using two fluorescent reporter proteins, green TagGFP2 and far-red Katushka, as described by Pereverzev et al. 2015. We have observed increased splicing-dependent NMD activity (Fig. 3E and 3F) and decreased 3’ UTR NMD activity (Fig. 3G and 3H) in mESCs treated with B2μM and C-L conditions, suggesting that Ca^2+^ exerts the opposite effect on splicing-dependent and 3’UTR-dependent NMD pathways in mESCs. We did not observe any change in the transcript levels of pluripotency and differentiation markers. This also raises the possibility that P-bodies, stress granule markers, and NMD in mESCs may play a major role in regulating markers other than pluripotency and differentiation through calcium signalling, and further studies are needed to identify those genes.

**Figure 5:**
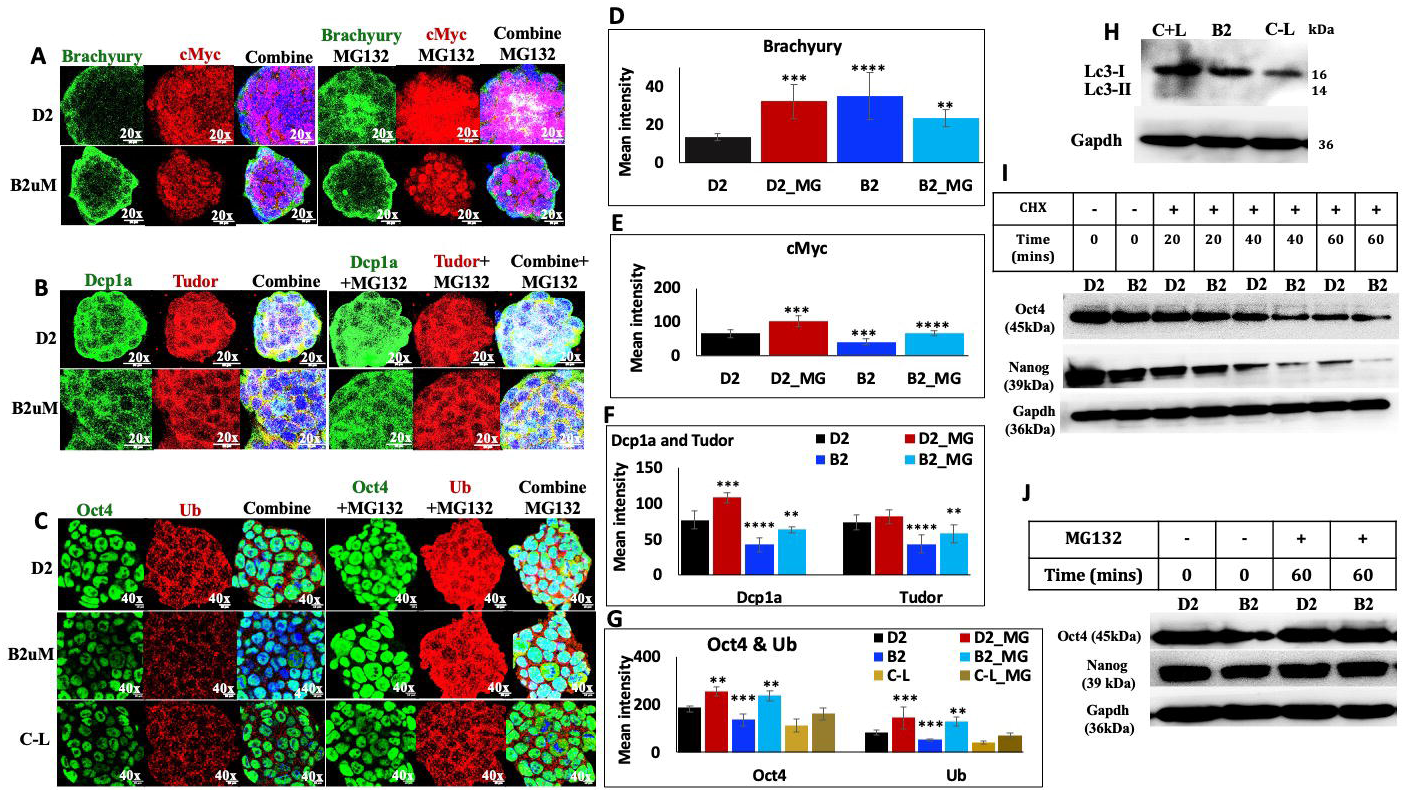
Role of calcium at post-translational level: Immunofluorescence (IF) detection of A) Brachyury, and cMyc, B) Dcp1a, and Tudor, C) Oct4 and Ub expression in mESCs cultured under different conditions (D2, B2μM and C-L) for 72 hours followed by MG132 (10μM) treatment for 1 hour (n=3). Figures 5A and 5B were taken at 20x magnification with a 20μm scale bar, and Figure 5C was taken at 40x magnification with a 10μm scale bar shown by representative images. IF experiments were done by thorough scanning the slide of different areas. The bar plot represents the mean intensity of the expression profile of D) Brachyury, E) cMyc, F) Dcp1a, and Tudor, G) Oct4 and Ub in mESCs colonies containing 20-50 cells from the confocal images (n=15) from different regions of the above experiments. H) Western blot detection of autophagy marker LC3 in mESCs cultured under different conditions (C+L, B2μM, C-L). (n=3) shown by representative images. Protein estimation was done using the Bradford method, and an equal amount of total protein from the cell lysate was loaded into the SDS-PAGE gel for all the samples. I) Cycloheximide (CHX) chase assay was performed for Oct4 and Nanog protein in mESCs cultured under D2, B2μM conditions for 72 hours, followed by the addition of CHX (50 ug/ml) and chased for every 20min for 1 hr 20 min using western blotting. (n=3) shown by representative images. Protein estimation was done using the Bradford method, and an equal amount of total protein for cell lysate was loaded into the SDS-PAGE gel for all the samples. J) Oct4 and Nanog expression was detected by western blot in mESCs cultured under D2, B2μM conditions for 72 hours, followed by the addition of MG132 (10uM) for 1 hour. (n=3) shown by representative images. Protein estimation was done using the Bradford method, and an equal amount of total protein from the cell lysate was loaded into the SDS-PAGE gel for all the samples. ***Note:*** *C+L= Control media containing Lif; C-L= control media without Lif; D2= C+L+ DMSO equivalent to B2*μ*M condition; D2_MG = D2+MG132; B2*μ*M= C+L+ BAPTA 2*μ*M; B2_MG = B2*μ*M+MG132; C-L_MG= C-L+MG132, n= biological replicates, BAPTA was used as calcium-modulating reagent*. *The mean intensity was calculated from the confocal images from different regions for all experiments using ImageJ software. An unpaired two-sample t-test was used to calculate the p-value using the MedCalc online statistical software calculator (MedCalc Software, 2024) with the mean and standard deviation. **p-value representation with asterisk <0.05 (), <0.01 (), <0.001 (), <0.0001 (****)***.

### Calcium doesn’t control the expression of pluripotency and differentiation markers at the post-transcriptional and translational level

We checked the global translation rate through polysome profiling and observed no change in the global translational rate upon Ca^2+^ manipulation with B2μM over 72 hours (Fig. 4A). However, we observed that the expression of the early translation initiation marker Eif1A was downregulated in B2μM-treated mESCs after 72 hours (Fig. 4B and 4C), which indicates a possibility of differential expression of translation markers upon Ca^2+^ manipulation. We have observed no change in the transcription of pluripotency and differentiation markers in control and B2μM-treated mESCs (S1_Supplementary Fig. 2A and 2B, Fig. 2I and 2J). We tested whether Ca^2+^ suppresses translation of pluripotency/differentiation markers by sequestering them in p-bodies/stress granules. To address this, we performed polysome profiling of mESCs treated with B2μM for 72 hours. We collected the cytoplasmic fraction (mRNP granules), 80S ribosomal subunit (80S), and polysome fraction. We assessed the expression of *Oct4, Nanog, Sox2,* and *Klf4,* as well as early mesoderm and ectoderm differentiation markers such as *Mixl1, Brachyury,* and *Pax6,* which showed high Ct values. We were unable to detect any significant difference in the transcript level of both pluripotency and differentiation markers in the cytoplasmic fraction of mRNP granules, the 80S and polysome fraction of control and B2μM-treated samples (Fig. 4D and 4E), indicating that pluripotency and differentiation markers are not held by p-bodies and might also not be regulated at the translational level. We observed no change in transcript levels of core pluripotency and early differentiation markers by qPCR upon BAPTA treatment (Figure 2I and 2J). To affirm this further, we blocked the transcription and translation of control, B2μM, C-L, C-L+T1nM treated mESCs after 72 hours with combined actinomycin-D (ACD) and cycloheximide (CHX) treatment for 2 hours. We checked the expression of pluripotency and mesodermal differentiation markers. We observed no significant change in their expression (Fig. 4F and 4G), illustrating that Ca^2+^ doesn’t regulate core pluripotency and differentiation markers at the transcriptional or post-transcriptional level. Taken together, our data signifies that calcium might regulate post-transcriptional processes but not pluripotency and differentiation markers through this route.

Undifferentiated ESCs either do not express or express low levels of differentiation markers (Ghimire et al., 2018). We noted that the transcript levels of early differentiation markers such as *Mixl1, Meox1* and *Brachyury* were expressed at CT values of 28±1, *Sox17* was mostly undetermined or expressed at a CT value of 34±1; *Pax6 and Sox1* were present at CT values of 26±1, in undifferentiated mESCs. These CT values indicate that differentiation-marker transcripts are present at low levels in undifferentiated mESCs. In addition, we detected transcripts of differentiation markers (*Mixl1, Brachyury*, and *Pax6*) in the polysome fraction of control and B2μM-treated mESCs (Fig. 4D and 4E), indicating that differentiation markers are being translated and might be expressed in undifferentiated mESCs or getting degraded. Since these mESCs showed a tendency to differentiate towards a mesoderm lineage (fibroblast-like morphology) spontaneously, we checked only Brachyury expression in the control, B2μM, and C-L conditions (Fig. 4H). We observed background Brachyury expression in undifferentiated mESCs and increased expression in B2μM and C-L conditions (Fig. 4H), indicating that calcium promotes Brachyury protein degradation in the C+L condition. Next, we performed MitoTracker-red staining and observed increased mitochondrial size and number after calcium reduction in B2μM and C-L conditions for 48 hours (Fig. 4I and 4J). We observed globular mitochondria in control mESCs and a network of branched mitochondria in B2μM and C-L in 48 hours (Fig. 4I), indicating that mitochondria undergo a morphological change from globular to tubular form upon Ca^2+^ reduction. Undifferentiated ESCs have globular, immature mitochondria and prefer glycolysis over oxidative phosphorylation. In contrast, during differentiation, mitochondria undergo a conformational change, and differentiated cells favour oxidative phosphorylation for energy production [Fu et al., 2019]. Thus, further studies are needed to examine the role of Ca^2+^in mitochondrial biogenesis and metabolic switching in mESCs.

### Calcium is important for the stability of pluripotency markers, p-body and stress granule markers, and degradation of differentiation markers

We also asked whether Ca^2+^ affects the post-translational degradation of proliferation, differentiation, core pluripotency markers, ubiquitination, P-body, and stress granule markers. Our data demonstrate an increase in the expression of cMyc, Dcp1a, Tudor, Oct4, and Ubiquitin (Ub) in both control and B2μM-treated mESCs upon MG132 treatment (Fig. 5 A, B, C, D, E, F, G), which suggests that Ca^2+^ levels might be important for the maintenance of homeostasis and stability of these proteins. We also observed increased Brachyury expression in the control condition after MG2132 treatment, which corroborates the increased expression of Brachyury upon B2μM treatment, but MG132 was unable to increase Brachyury protein expression in B2μM-treated mESCs, but rather it led to its decreased expression (Fig. 5A and 5D). Earlier findings suggest that proteasomal inhibition can lead to protein degradation through activation of other degradation pathways, such as autophagy (Wang et al., 2019). Furthermore, we noted that MG132 treatment rescued only partial protein levels of Dcp1a and Tudor in B2μM-treated mESCs compared with control mESCs (Fig. 5B and 5F), which also implies the role of other signalling pathways in regulating P-body and stress granule markers upon Ca^2+^ reduction. Interestingly, we also noted a decrease in ubiquitin (Ub) expression using anti-ubiquitin antibody (detect ubiquitin, polyubiquitin, and ubiquitinated proteins), upon B2μM treatment and its upregulation with MG132 treatment through IF (Fig. 5C and 5D), suggesting that upon Ca^2+^ reduction, degradation of pluripotency markers might be independent of the ubiquitin-proteasomal system. Several studies have suggested that MG132 increases the expression of ubiquitinated proteins using an anti-ubiquitin antibody (Liang et al., 2025; Meiners et al., 2002). In addition, our phosphoproteomic data detected significant decreased expression of E3 ubiquitin protein ligases Ubr4, Trip12, Stub1, and Rbbp6, as well as increased expression of E3 ubiquitin protein ligase Nedd4 upon BAPTA treatment (Fig. 7A and Supplementary file S2 (S2)_Supplementary Table 1, 2, 3, 5), which indicates that calcium might be playing a role in the degradation of proteins through the proteasomal degradation pathway in undifferentiated mESCs. Next, we observed very low LC3-I expression and a faint LC3-II band in undifferentiated mESCs, which decreased further under B2μM and C-L conditions, suggesting autophagy might not be playing a role in the degradation of pluripotency and differentiation markers upon Ca^2+^ reduction (Fig. 5H). We also could not detect the ATG7 autophagy marker on the Western blot. Several studies have shown that autophagosomes can be detected by analysing diffuse cytoplasmic LC3-I and punctate LC3-II (which correlates with autophagosomes) by IF (Holt et al., 2011; Rosenfeldt et al., 2012; Mizushima et al., 2010). We detected diffuse and punctate LC3, which decreased under B2μM and C-L conditions (S1_Supplementary Fig. 6A and 6B). Further, we were unable to detect key autophagy phosphoproteins, including p62, Ulk1, Beclin-1, Atg 13, Fip 200, and Atg 4B (Licheva et al., 2022, Eickhorst et al., 2024, Matsumoto et al., 2011) in our phosphoproteomic dataset (Supplementary file S2 (S2)_Supplementary Table 1), which indicates that autophagy-related phosphoproteins might be present in low amounts in mESCs or autophagy might be active at a low level in mESCs. In addition, our study indicates that the ubiquitin-proteasome system might be important for the degradation of differentiation markers in undifferentiated mESCs, and that its low expression upon Ca^2+^reduction might be important for Brachyury stability and for activating other signalling pathways, apart from autophagy, to determine lineage specificity. Further studies are needed to define the role of Ca^2+^ signalling in controlling ubiquitination-mediated proteasomal degradation for the regulation of differentiation markers in mESCs. We have further checked the stability of Oct4 and Nanog using a CHX chase assay as described by Kao et al., 2015. We chased the expression of Oct4 and Nanog for 1hr 20 min with a 20 min time interval and observed that their degradation was faster in B2μM-treated mESCs, indicating that Ca^2+^ is important for the maintenance of the stability of core pluripotency markers (Fig. 5I). Furthermore, we observed an increase in the level of Oct4 and Nanog proteins in B2μM-treated mESCs with MG132 treatment (Fig. 5J), suggesting that Ca^2+^ maintains the stability of pluripotency markers at the post-translational level.

**Figure 6:**
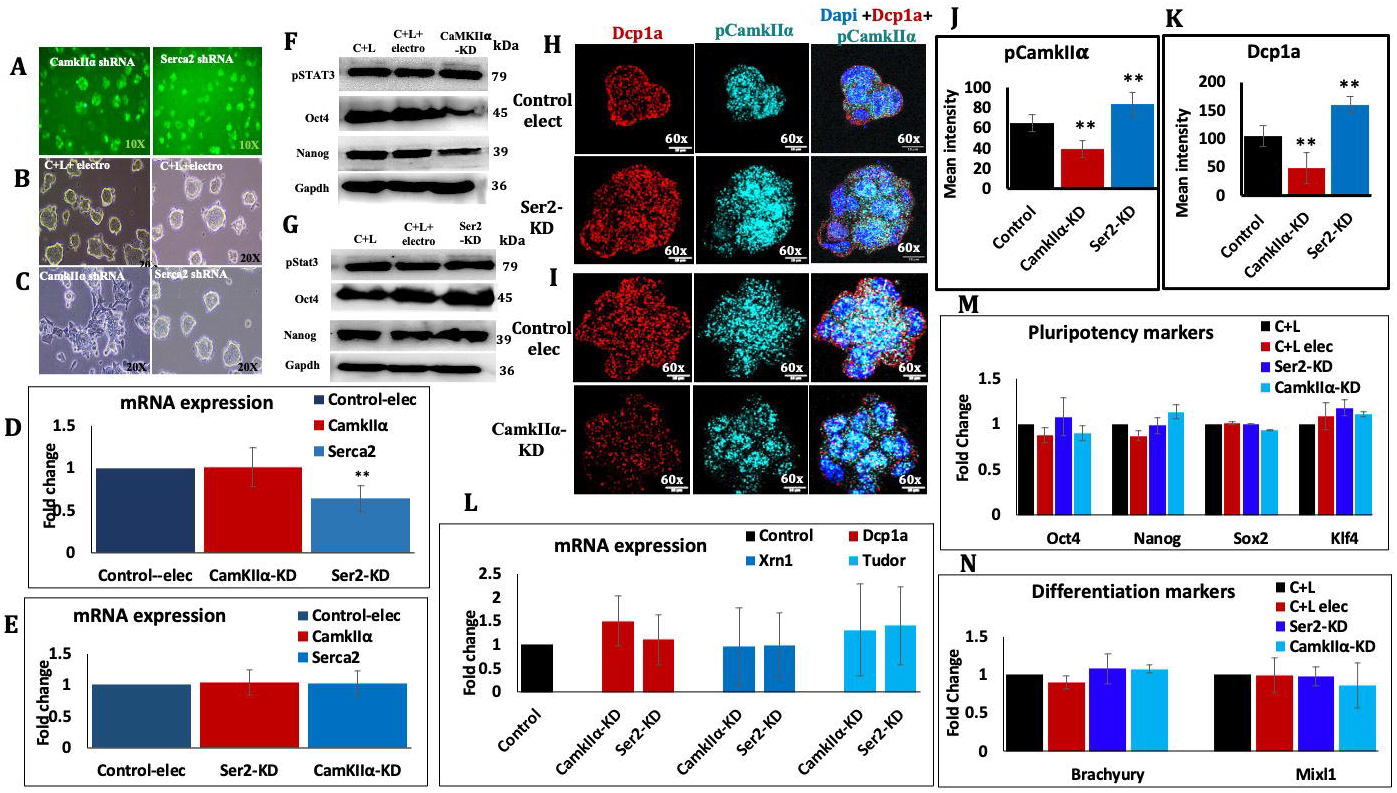
knockdown of pCamkIIα in mESCs. **A)** Detection of GFP expression of *CamkIIα* shRNA and *Serca2* shRNA transfected by the electroporation method in mESCs using fluorescent microscopy at 10X magnification (n=3) shown by representative images. Brightfield images showing the morphology of the **B)** control mESCs, **C)** *CamkIIα* shRNA (CamkIIα-KD) and Serca2 shRNA (Ser2-KD) transfected mESCs at 48 hours (n=3). **D)** QPCR of *CamkIIα* and *Serca2* expression in the mESCs at 48 hours upon CamkIIα-KD and Ser2-KD, respectively, n=3. **E)** QPCR of *CamkIIα* expression in the Ser2-KD mESCs and *Serca2* expression in the CamkIIα-KD mESCs at 48 hours (n=3). **F, G)** Western blot detection of Oct4, Nanog, pStat3, and Gapdh expression in the CamkIIα-KD and Ser2-KD mESCs at 48 hours (n=3) shown by representative images. Protein estimation was done using the Bradford method, and an equal amount of total protein from the cell lysate was loaded into the gel for all samples. **H, I)** IF detection of Dcp1a and pCamkIIα co-expression in CamkIIα and Ser2-KD mESCs at 48 hours at 60x magnification with a 10 μm scale bar shown by representative images. (n=3). IF experiments were done by thorough scanning the slide of different areas. **J, K)** Bar plot represents the mean intensity of pCamkIIα and Dcp1a expression in CamkIIα-KD and Ser2-KD mESCs colonies containing approximately 7-20 cells, calculated from the confocal images (n=15) of the above experiments. **L)** QPCR of p-bodies and stress granule markers in the mESCs at 48 hours upon pCamkIIα-KD and Ser2-KD. (n=4). **M, N)** QPCR of pluripotency and differentiation markers in the mESCs at 48 hours upon CamkIIα-KD and Ser2-KD. (n=3). ***Note:*** *CamkIIα shRNA (CamkIIα-KD) and Serca2 shRNA (Ser2-KD), control elec = control shRNA, n=biological replicates* ***Note:** The mean intensity was calculated from the confocal images from different regions for all experiments using ImageJ software.* An unpaired two-sample t-test was used to calculate the p-value using the MedCalc online statistical software calculator (MedCalc Software, 2024) with the mean and standard deviation. ***p-value representation with asterisk <0.05 (*),*** *<0.01 (**), <0.001 (***), <0.0001 (****)*.

**Figure 7:**
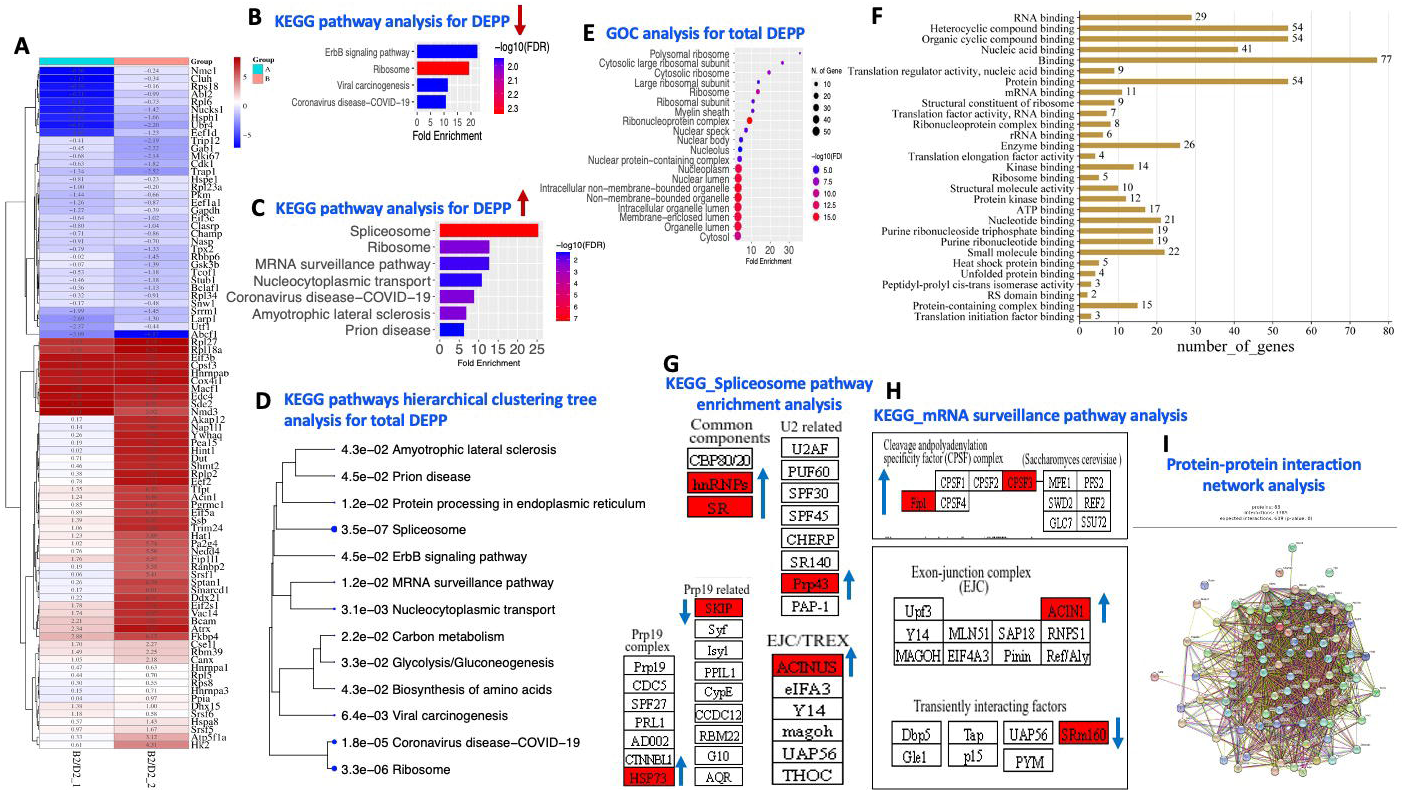
Calcium and phosphoproteomics in mESCs: **A)** Heat map shows log2fold change of DEPP in B2μM treated mESCs compared to D2 (control) mESCs detected by mass spectrometry (n=2, each set pooled phosphoproteomics). **B, C, E)** KEGG pathway and GOC enrichment analysis were done using ShinyGO 0.80 software for the DEPP represented in the heat map (A). **D)** The hierarchical clustering tree shows the correlation among the most significant pathways found in KEGG pathway enrichment analysis for all DEPP. **F)** The bar plot shows the string enrichment function analysis of all the DEPP demonstrated in the heat map. **G, H)** KEGG pathway enrichment analysis was done for spliceosome and mRNA surveillance pathways, all the DEPP represented in the heat map (A). I) String PPI network analysis was performed for all DEPP represented in a heat map (A). **Note:** Arrows represent the upregulated and downregulated genes ***Note:*** *C+L = Control media containing Lif; B2*μ*M = C+L + BAPTA 2*μ*M; D2 = C+L+ DMSO equivalent to B2μM condition, BAPTA was used as calcium-modulating reagents. p-value was calculated using the one-way Chi-square test. **p-value representation with asterisk <0.05 (*), <0.01 (**), <0.001 (***), <0.0001 (****)***.

### Calcium maintains the stability of Oct4, Nanog, and Dcp1a by a pCamkII alpha-dependent mechanism

To confirm the role of calcium signalling in regulating pluripotency and p-body markers, we knocked down CamkIIα, a major downstream target of calcium signalling, to assess the effects of decreased calcium levels and Serca2 knockdown to observe the effect of increased calcium on calcium signalling. We used the electroporation method to deliver CaMKIIα shRNA to silence pCamkIIα expression. We noted that 80-90% of mESCs expressed *CamkIIα* shRNA, as shown by their GFP expression (Fig. 6A). Moreover, we observed that CaMKIIα-KD mESCs exhibited loose dome-shaped colonies with fibroblast-like cells emerging from their edges (Fig. 6B and 6C), which was similar to the B2μM-treated mESCs (Fig. 2K). The *CamkIIα* shRNA targets and silences the *CamkIIα* mRNA and prevents the production of unphosphorylated and active pCamkIIα protein (Moore et al., 2010; Hirst et al., 2022). We were more interested in exploring the role of the active form of CamkIIα and thus focused our study on pCamkIIα and checked the expression of pCamkIIα upon CamkIIα-KD. We were unable to detect pCamkIIα by Western blot, either due to no signal or a noisy signal with ECL Amersham GE kit, so we performed thorough IF scanning for pCamkIIα upon CamkIIα-KD. However, we were unable to detect any significant decrease in the level of *CamkIIα* mRNA but observed a significant decrease in the level of pCamkIIα protein (pCamkIIα-KD) upon *CamkIIα* shRNA treatment (CamkIIα-KD) (Fig. 6D and 6I), which suggests the possibility that *CamkIIα* shRNA might have exerted an indirect mechanism in regulating pCamkIIα protein expression rather than canonical knockdown efficiency, and this might be due to a decrease in RNA pool was not detected by qPCR or through translational repression, which is a another mechanism action of small RNAs including siRNA and miRNAs, where they trigger translation repression without or with minimally affecting the stabilization of mRNAs (Ma et al., 2013, Carthew and Sontheimer 2009, Li et al., 2021). Furthermore, we detected a significant decrease in the expression of Oct4 and Nanog proteins upon pCamkIIα-KD by western blotting (Fig. 6F), similar to B2μM-treated mESCs (Figure 2A, 2B, 2C, 2E). Several studies have demonstrated a decrease in the expression of Oct4, Nanog, Sox2 and cMyc upon inhibiting CamkII phosphorylation or CamkII or CamkIIγ in cancer cells and cancer stem cells (Shin et al., 2019, Chai et al., 2015, Han et al., 2015). However, we did not observe any change in the transcript levels of either pluripotency or differentiation markers (Fig. 6M and 6N), as in B2μM-treated mESCs (Figure 2I and 2J).

In addition, we observed that the pStat3 protein level was unaffected in pCamkIIα-KD mESCs, which again signifies that Ca^2+^ might not be playing a role through the Jak-Stat pathway (Fig. 6F). We further observed a decrease in the expression of the P-body marker Dcp1a upon pCamkIIα-KD (Fig. 6I), which indicates that Ca^2+^ is important for the stability of Dcp1a through a pCamkIIα-dependent mechanism.

We also performed Serca2 knockdown (Ser2-KD) using shRNA against highly expressed Serca2 calcium pump, as noted in our QPCR data. We observed a decrease in the mRNA pool of the *Serca2* upon Ser2-KD (Fig. 6D). We noted that Ser2-KD mESCs exhibited compact colonies as compared to control mESCs (Fig. 6B and 6C), which was similar to T1nM-treated mESCs (Fig. 2K). Furthermore, we observed an increase in the expression of Oct4 and Nanog proteins upon Ser2-KD (Fig. 6G),which was similar to T1nM-treated mESCs (Fig. 2A, 2B, 2C, 2F, 2I, and 2J). We observed no change in the transcript levels of pluripotency and differentiation markers (Fig. 6M and 6N) or in the protein level of pStat3 in Ser2-KD mESCs (Fig. 6G), again suggesting that calcium might not act through the Jak-Stat pathway. Furthermore, we observed increased expression of Dcp1a and pCamkIIα upon Ser2-KD (Fig. 6H), indicating that cytosolic Ca^2+^ plays a role in Dcp1a biogenesis through pCamkIIα.

### Identification of phosphoproteins altered upon calcium manipulation using a phosphoproteomics approach

We conducted a pilot-scale exploratory analysis of pooled phosphoproteomic profiles from control and B2μM-treated mESCs (n=2 per set, pooled phosphoproteomics). We identified 53 significantly upregulated phosphoproteins (PPs) and 35 downregulated PPs in B2 samples, as shown by the heatmap (Fig. 7A and Supplementary file S2_Supplementary Table 1, 2, 3, 5). Using Proteome Discoverer Software (Thermo Fisher Scientific), we identified phosphosites in the phosphopeptides from mass spectrometry (MS) data. We observed mainly serine-threonine phosphosites, along with a few tyrosine sites (Supplementary file S2_Supplementary Tables 1 and 4), indicating that Ca^2+^ manipulation through BAPTA primarily influences a broad category of serine-threonine PP. Using ShinyGO 0.80 software, we performed KEGG pathway, tree plot, GO cellular component (GOC), and STRING function enrichment analyses for the differentially expressed phospho-proteins (DEPP) shown in the heat map (Fig. 7B, C, D, E, F). We observed that spliceosome and mRNA surveillance PP were highly enriched in our dataset and performed further KEGG pathway enrichment analysis (Fig. 7G and 7H). We found that our PP are part of these complexes labelled in red. We further observed that DEPP exhibits 1385 protein-protein interactions (PPI), with an expected 689 PPI (p=0), using STRING PPI network analysis (Fig. 7I). Next, we identified DEPP-related pathways using STRING bio process enrichment analysis (SBPEA). We conducted GO biological function (GOB), GO molecular function (GOF) and GOC analysis for DEPP related to metabolism listed in the legend of Supplementary Fig. 4A, and observed that they are positive and negative regulator of metabolism, part of the mRNP complex and displayed nuclear and organelle localisation, which suggests a widespread functional role of these proteins (S1_Supplementary Fig. S7A, 7B, 7C). We also performed the GOC analysis of a few DEPP, including Hk2, Hspa8, Atp5a1, Cox4i1, Pkm, and Gapdh, identified for ATP metabolism. We observed mitochondrial localisation (S1_Supplementary Fig. S7J), which supports their role in mitochondrial biogenesis and oxidative phosphorylation. We also noted that several PP related to translational machinery including Rps18, Rp16, Eif3c, Abcf1, Eef1a1, Eef2, Eif5a, Rpl18a, Eif2s1, Rpl5, Rplp2, Rpl34, Eef1d, Rpl27, Eif3b, Rpl23a, Rps8, and Larp1 were highly differentially regulated (Fig. 7A), despite global translation rate was unaffected upon Ca^2+^ reduction and conducted GOC, GOB network and GOF analysis for them (S1_Supplementary Fig. S7G, 7H and 7I). We observed that these proteins are also part of the mRNP complex and have a role in the macromolecular biosynthetic pathway, which exemplifies their other role apart from controlling the translation rate. Next, we performed GOB and GOF analyses for DEPP implicated in nucleosome and chromatin assembly, including Smarcd1, Nasp, Hat1, Atrx, and Nap1l1 (S1_Supplementary Fig. S7D and 7E). We observed that they have chromatin– and histone-binding properties and play a role in DNA replication-independent chromatin assembly. Further, we conducted GOB network analysis of DEPP, including CSe1, smarcd1, Hk2, Pkm, Tfpt, Tcof1, champ1, nasp, Macf1, Hat1, Bcam, Atrx, Nmd3, Sde2, Ubr4, Hsph1, Trap1, nucks, Cdk1, Mki67, Tpx2, champ1 (S1_Supplementary Fig. 7F). We observed that they exhibit cross-talk in the kinetochore process, chromatin condensation, nuclear division, mitotic cell cycle, and chromatin remodelling. GOB network analysis also identified DEPP, including Cdk1, Mki67, Nucks1, and Tpx2, that are crucial for mitotic cell division and mitochondrial ATP synthesis, implicating their role in G2/M cell cycle arrest and oxidative phosphorylation (S1_Supplementary Fig. S8A and 8B). However, further studies are needed to confirm the role of Ca^2+^in the regulation of the above processes in mESCs.

## Discussion

In recent years, pluripotent stem cells have gained significant importance in regenerative medicine for studying cell development and differentiation. However, the role of Ca^2+^ signalling in maintaining pluripotency remains unexplored. Previous work in Drosophila, mesenchymal stem cells, and mESCs has shown that stem cells differentiate upon Ca^2+^reduction (Shim et al., 2013; Vallet et al., 2025; MacDougall et al., 2025; Zhu and Zhang, 2019; Apáti et al., 2016). Our study showed that low calcium induces spontaneous differentiation of mESCs, reflecting a broadly conserved phenomenon in stem/progenitor cells across species. In stem cell biology, there is a persistent effort to understand the mechanism underlying ESC pluripotency. Nevertheless, the full repertoire of molecules that control the ESC state is not yet fully understood. Several studies have reported heterogeneous NANOG expression in mESCs, but the mechanism underlying it remains unclear (Torres-Padilla and Chambers 2014). Previous findings in hESCs have also indicated heterogeneous expression of Ca^2+^ signalling (Huang et al., 2017). Our data indicate a correlation between heterogeneous Oct4 and Nanog expression and heterogeneous Ca^2+^ and pCamkIIα levels in mESCs, suggesting that calcium might be related to heterogeneous pluripotency-marker expression.

Earlier reports suggest that activation of the calcium receptor Ryr2, the calcium-sensing receptor (CaSR), and increased calcium levels promote differentiation of human and mouse ESCs toward cardiac differentiation (Meng et al., 2025; Sun et al., 2013). Interestingly, we noted that upon Ca^2+^reduction, mESCs lose pluripotency and show a tendency to acquire mesoderm-like lineage, suggesting that Ca^2+^signalling might be important in undifferentiated mESCs to mask the expression of mesoderm markers. However, additional experiments with protein markers, comprehensive gene expression profiling or differentiation assays are needed to confirm the role of Ca^2+^signalling in mediating mesoderm differentiation of mESCs. Furthermore, our data indicate that high Ca^2+^ mediates Brachyury degradation, while low calcium and a low ubiquitin might be important for its stability by preventing its degradation from UPS; this also suggests activation of other signalling pathways upon calcium reduction to determine lineage specificity. Moreover, our data suggest that calcium may not control the Lif-Jak-Stat3 pathway for regulating core pluripotency and differentiation markers at the transcriptional level, but instead controls the stability of these fundamental regulators at the post-translational level.

Our pCamkIIα KD data indicate that pCamkIIα might be important for regulating the expression of Oct4 and Nanog proteins in mESCs. We have also observed co-localisation of Oct4 and pCamkIIα, but have not checked whether and how pCamkIIα phosphorylates the core pluripotency markers Oct4 and Nanog, or how this results in their downregulation. pCamkIIα is a serine-threonine kinase involved in phosphorylating a wide array of proteins (Clapham 2007), suggesting it may regulate the phosphorylation status of these core pluripotency markers. Earlier studies have shown that phosphorylation of Oct4 and Nanog at serine residues decreases their stability and directs them toward ubiquitination-mediated protein degradation in mESCs (Bae et al., 2017; Kim et al., 2014). However, our study demonstrates increased expression of Oct4 and Nanog in the presence of high calcium and pCamkIIα. Moreover, our data also indicate that Ca^2+^ may regulate Oct4 and Nanog expression independently of the ubiquitination-mediated proteasomal degradation upon calcium reduction. In addition, we were unable to detect phosphorylated Oct4 and Nanog in our phosphoproteomic dataset, which indicates that the phosphorylated form of Oct4 and Nanog might be present at a low level in mESCs in the presence of high and low Ca^2^, which suggests that other mechanisms might also be playing a role in the regulation of expression of Oct4 and Nanog through calcium signalling. Several reports suggest that protein degradation can occur by 20S-or 26S-mediated proteolysis driven by structural disorder, independently of polyubiquitin tagging or without the 19S proteasome subunit (Shang and Taylor 2011; Ben-Nissan and Sharon 2014). Previous findings have advocated that Ca^2+^ and Ca^2+^-binding proteins promote conformational change upon binding to target proteins and modulate their activity and stability (Clapham, 2007, Strynadka, and James, 1991, Hołubowicz et al., 2017) Therefore, further conformational, structural, biochemical, and molecular studies are needed to understand the in-depth mechanism of how Ca^2+^ regulate the activity and stability of pluripotency markers using targeted experiments such as co-immunoprecipitation, in vitro phosphorylation, site-directed mutagenesis, phosphomimetic/mutant variants of Oct4 and/or Nanog, rescue with CamkIIα expression or FRET (Fluorescence Resonance Energy Transfer), CD Spectroscopy (Circular Dichroism), cryo-EM, computational and in-silico modelling.

Our phosphoproteomic data has identified several RNA-binding and ribosomal PP including Edc4, Pa2g4, Nmd3, Ddx21, Srsf5, Srsf6, Srsf1, EDC4, Fkbp4, Cpsf3, Fip1l1, Hnrnpab, Hnrnpa, Hnrnpa3, Rbm39, Dhx15, Acin1, Snw1, Larp, Srrm1, Clasrp, Rps18, Rars, Eif3c, Eef1a1, Eif5a, Rpl18a, Eif2s1, Rpl5, Rps8, Rplp2, Rpl34, Eef1d, Rpl27, Eif3b, Rpl23a implicated in mRNP complex formation, alternative splicing, alternative polyadenylation, mRNA stability, 5′TOP mRNAs stability, miRNA biogenesis, mRNA decapping and NMD through GOB analysis and literature review (Martinez et al., 2013, Hochstoeger et al., 2024, Chiang et al., 2013, Decker and Parker, 2012), which suggests a widespread role of Ca^2+^at a post-transcriptional level rather than global translational control. Our data also show that Ca^2+^ affects Dcp1a, Xrn1, Tudor, and NMD reporters. Yet, it remains unclear what role they play in the regulation of pluripotency and cell fate determination.

Furthermore, we did not observe any key phospho-apoptosis markers, including Bcl-xL, c-jun, H2AX, phospho-histone H3, and FAS (Stokes et al., 2012; Solier and Pommier, 2009; Park and Kim, 2012; Holmstrom and Eriksson, 2000), in our phosphoproteomic dataset. Indeed, our phosphoproteomic data detected the apoptosis marker BCL-2-associated transcription factor 1, but we did not observe any significant change in its protein profile (Supplementary file S2_Supplementary Table 1), which indirectly supports that apoptosis might not be dysregulated. In addition, we did not observe any ER stress-related key phosphoproteins such as Perk, eIF2α, and Ire1α (Chen et al., 2023) in our phosphoprotein dataset (Supplementary file S2_Supplementary Table 1). Moreover, ER stress triggers the heat shock protein response (Liu and Chang, 2008). We have detected several heat shock phosphoproteins, such as Hsp-90 alpha, Hsp 90 beta, Hspa8, Hspa4, Hspb1, Ahsa1, Hsph1, and Carhsp1 in our phosphoproteomic dataset (Supplementary file S2_Supplementary Table 1). Among these, only Hspa8 and Hsph1 were significantly upregulated and downregulated, respectively, indicating that ER stress might not be substantially altered.

Taken together, our data indicate that calcium might have a broad role in modulating various biological processes such as cell cycle arrest, protein stability, RNA splicing, mRNA stability, ribosome biogenesis, mRNP granule formation, ubiquitination, metabolism, telomere maintenance, oxidative phosphorylation, mitochondrial biogenesis, chromatin regulation, and RNA polyadenylation for the maintenance of mESCs identity.

## Materials and Methods

### Culturing of mESCs and calcium modulation

This study used mESCs E14TG2a and E14 cell lines. The work was mainly conducted using E14 mESCs. E14TG2a cell line was obtained from Dr Ramkumar Sambasivan lab (Former professor at inStem, Bangalore, India 2015) and E14 mESCs were obtained from the Dr Dasaradhi Palakodeti Lab (inStem, Bangalore, India). These cell lines were maintained on 0.1% gelatin at 37 °C in a 5% CO2 incubator in ES medium containing high-glucose DMEM (GibcoTM), ESGRO recombinant mouse LIF, 15% ES cell FBS (GibcoTM), β-Mercaptoethanol (GibcoTM), 1x NEAA (GibcoTM), and gentamicin (GibcoTM). To induce spontaneous differentiation, mESCs were trypsinised with 0.05% Trypsin-EDTA (GibcoTM) and cultured in ES media without LIF. For BAPTA-AM (InvitrogenTM) and thapsigargin treatment, mESCs were cultured in the presence of LIF with different concentrations of BAPTA (1 μM and 2 μM) and thapsigargin (250 pM, 500 pM, and 1nM) for 72 hours. BAPTA and thapsigargin stock solutions were prepared using dimethyl sulfoxide (DMSO). An equal amount of DMSO was added to the control media as compared to BAPTA and thapsigargin. The medium was changed every day. All experiments were done at 72-hour intervals, except mitochondrial staining, which was done at 48 hours, when mitochondrial morphological changes were detected. The transcript profile was studied at 48 hours and 4 days to determine whether transcripts remained unchanged after calcium manipulation at shorter and longer time intervals. Cell cycle, proliferation and c-Myc staining were studied for 5 days with 72-hour BAPTA treatment and a 48-hour rescue condition.

### Cell proliferation assay and cell cycle assay

mESCs proliferation was calculated using the trypan blue method, and cell cycle analysis was done by PI staining. Briefly, mESCs were fixed in 70% ethanol and stored at –20 °C. Fixed cells were centrifuged at 1000 rpm for 3-5 min and washed with PBS 2-3 times and PI staining solution (80 μg/ml PI, 100 μg/ml RNase A and 0.1 % Triton X-100) for 7 hrs at 4 °C. After incubation, mESCs were washed with PBS 2-3 times and cell cycle analysis was done using flow cytometry and BD software.

### MitoTracker Red staining, Fluo-4 staining, LysoTracker Staining, and Immunofluorescence staining

Cells were grown on coverslips in ES media containing BAPTA (2 μM) and thapsigargin (1nM). After 72 hours, cells were incubated with media containing Fluo-4 (1 μM), LysoTracker (100 nM), and MitoTracker Red (100 nM) for 45 minutes, followed by 2-3 washes with media, then immediately taken for live-cell imaging. For Fluo 4 staining, cells were incubated at 37 °C in a 5% CO2 incubator for 15 minutes before live-cell imaging. Imaging was performed using a confocal microscope at 40x magnification. For immunostaining, cells were grown on coverslips and treated with BAPTA (2 μM) and thapsigargin (1 nM) for 72 hours, then fixed in 4% paraformaldehyde (PFA) overnight at 4 °C. After incubation, PFA was removed, and cells were washed twice with PBS. The fixed cells were permeabilised using 0.4% Triton X-100 for 2 hours at room temperature (RT). The cells were then blocked with blocking buffer (5% BSA and 0.1% Tween-20 in PBS) for 2 hrs at RT. After blocking, cells were incubated with primary antibodies at 4 °C overnight or 3 hours at RT, followed by two washes with PBS for 5 minutes each. The primary antibodies used were Oct3/4 (Invitrogen), Nanog (Invitrogen), Eif1a (Abcam), Dcp1a (Sigma), Xrn1, beta-actin (mRNP) (Santa Cruz), pCamkIIα (Cell Signalling Technology), pStat3 (Abcam), Ubiquitin (Abcam), cMyc (Abcam), and Brachyury (Santa Cruz). Further, secondary antibodies (Jackson ImmunoResearch Laboratories (West Grove, PA) were added and incubated for 1-2 hr at RT. After incubation, cells were washed twice with PBS for 5 minutes and mounted on a glass slide using mounting medium containing DAPI. For IF experiments, confocal images were used to calculate mean intensity in ImageJ software for gene expression analysis.

### Gene expression analysis

RNA was extracted using the TRIzol (Invitrogen, Carlsbad, CA, USA) method. Briefly, cells were lysed and homogenised using TRIzol. Then, 1/5 volume of chloroform was added to the TRIzol, and samples were centrifuged at 12000 rpm at 4 °C for 20 min. After centrifugation, the aqueous layer was collected, an equal volume of isopropanol was added, and samples were incubated at –80 °C overnight. After incubation, samples were centrifuged to precipitate RNA. After centrifugation, precipitated RNA was washed with 70% ethanol, dried, and dissolved in nuclease-free water and stored at –80 °C. Further, 1 μg of RNA was reverse transcribed with Superscript II or Superscript III reverse transcriptase (Invitrogen) according to the manufacturer’s protocol. Real-time polymerase chain reaction (PCR) was performed using AB Invitrogen with Power Sybr Green (Thermo Fisher Scientific) according to the manufacturer’s instructions. The gene expression analysis of pluripotency and differentiation markers was calculated using the formula 2^∧^ΔΔct and normalised to glyceraldehyde 3-phosphate (Gapdh).

### Protein extraction and western blotting

For protein extraction, mESCs were lysed using radioimmunoprecipitation assay (RIPA)buffer. The samples were then centrifuged at 12000 rpm for 10 minutes at 4 °C. The supernatant was collected, and protein estimation was quantified by the Bradford method using BSA as a standard. An equal amount of protein (5-10 μg) was separated on a 12% SDS-PAGE gel. PageRuler Prestained Protein Ladder with range 10kDa-180kDa and 10kDa-250kDa were used as markers (Thermo Scientific™). Proteins were electrophoretically transferred into polyvinylidene difluoride membranes (Bio-Rad) for western blot analysis. The blots were blocked with blocking buffer containing 5% BSA in TBST (20 mM Tris pH 7.5, 150 mM NaCl, and 0.1% Tween-20) for 1-3 hours. The blots were cut horizontally at specific molecular weights (M.W.) and incubated with primary antibody at 4°C overnight or for 3-4 hours at RT. After incubation, blots were washed twice with TBST, and membranes were incubated with horseradish peroxidase (HRP)-conjugated secondary antibody for 2 hrs. Signals were detected with ECL substrate (Amersham GE) using ImageQuant. Primary antibodies included Oct3/4 (Invitrogen), Nanog (Invitrogen), Gapdh (Invitrogen), cMyc (Abcam), and pStat3 (Abcam); secondary antibodies were anti-rabbit HRP and anti-goat HRP. Due to the similar M.W. of Gapdh (36kDa), Oct4 (45kDa), and Nanog (39kDa), all proteins were run separately on the SDS-PAGE gel. Gapdh was used as a loading control.

### Cycloheximide chase assay, MG132 treatment for analysing protein stability, and actinomycin D and CHX assay for post-transcriptional activity detection

mESCs were cultured in the absence and presence of B 2 μM for 72 h, followed by treatment with 50 µg/mL CHX for 1 h, and Oct4 and Nanog protein levels were detected every 20 min by Western blot. For the MG132 experiment, mESCs were cultured in the absence and presence of BAPTA (2 µM) for 72 hours, followed by 10 µM MG132 treatment for 1 h, and analysed by Western blot and IF. For posttranscriptional analysis of pluripotency markers, mESCs were treated with 5ug/ml of actinomycin-D (ACD) and CHX (50ug/ml) for 2 hr after culturing the mESCs in C+L, B2μM, T1nM, C-L, C-L+T1nM, and samples were extracted in Trizol for RNA purification.

### Electroporation

Electroporation was performed using an Invitrogen Neon Transfection System (100 µL) according to the manufacturer’s instructions. Briefly, mESCs were trypsinised, washed with PBS, and centrifuged at 1000rpm for 3-5 min. The mESCs pellet was then resuspended in the resuspension buffer (Buffer R) containing the required plasmid (*CamkIIα* shRNA, *Serca2* shRNA, Design 1 Endoplasmic reticulum (D1ER) calcium sensor for measuring endoplasmic reticulum calcium as described by Waldeck-Weiermair et al. (2015), and NMD reporter plasmids [NMD reporter plasmids were kind from Konstantin A. Lukyanov from Sergey A. Lukyanov lab, Russian Academy of Sciences, Russia]). Electroporation was performed at 900 V, 3 pulses, and 23 ms pulse width. After electroporation, mESCs were immediately transferred to ES media without antibiotics and incubated at 37 °C in a 5% CO2 incubator. After 24 hours, fresh media containing antibiotics was added, and GFP expression was analysed between 48 and 72 hours. After 48 hours, mESCs were collected in RIPA buffer for western blot, TRIzol for RNA extraction and fixed in 4% for immunofluorescence staining.

### Polysome profiling

mESCs were cultured in the presence of B2μM for 72hour and pellet was resuspended in polysome lysis buffer (20mM Tris pH-7.4, 200mM KCL, 5mM Mgcl2, 0.7% Triton X-100, 1x protease inhibitor, RnaseoOUT, 100ug/ml CHX, 1mM DTT). A 15-45% sucrose gradient was prepared in gradient buffer (20 mM Tris, pH 7.4, 5 mM MgCl2, 100 mM KCl, 1 mM DTT, 100 μg/ml CHX) using a gradient mixer. An equal amount of RNA/260 OD was loaded into the gradient and centrifuged at 39000 rpm at 4 °C for 2.5 hr. The polysome peak was analysed using a polysome profiler, and cytoplasmic mRNP granule, 80s ribosome, and polysome fractions were collected in absolute ethanol. Samples were stored at –80 °C overnight, and RNA was isolated by collecting pellets from ethanol fractions in TRIzol or using an RNA extraction kit (Qiagen). cDNA synthesis was carried out using Superscript II or Superscript III according to the manufacturer’s instructions.

### Phosphoproteomics

We conducted a pooled phosphoproteomics experiment in n=2 replicates. mESCs were collected from one 60mm dish and one 35mm dish, each dish cultured on different days and pooled to form one set. mESCs were cultured for 72 hours in the presence of B 2μM, and proteins were extracted using RIPA buffer containing protease and phosphatase inhibitors (Roche). Proteins were then precipitated with 15% TCA, followed by a 90% acetone wash. The protein pellet was dissolved in 8% urea buffer (8% urea in 100 mM ammonium bicarbonate buffer) containing 5 mM DTT and 1× phospho-stop inhibitor (Roche), then incubated at RT for 35 min to denature the proteins fully. Protein levels were estimated using the Bradford method. Equal amounts of protein (500mg and 200mg) were taken from control and BAPTA-treated samples and incubated at 56 °C for 45 min with 5 mM DTT. Samples were further treated with 10 mM iodoacetamide and incubated at 56 °C for 45 min. After incubation, Samples were diluted 10-times with ammonium bicarbonate buffer and incubated at 37 °C overnight with trypsin gold (mass spec grade, Promega) in a 1:100 ratio. After incubation, peptides were collected using 1% FA. Alternatively, during protocol standardisation, the remaining protein pellet after the first digestion was dissolved twice in ammonium bicarbonate buffer to reactivate trypsin gold, supplemented with 500 ng trypsin gold, and incubated at 37 °C for 8-9 h and overnight, respectively; peptides were collected using 1% formic acid. Peptide concentrations were monitored on a NanoDrop by measuring 180-300 nm UV spectra and the peak around 190-220 nm; peptides were quantified using the A205 preprogrammed absorbance application (NanoDrop One Spectrophotometer, Thermo Fisher Scientific). Peptides were purified using an Oasis HLB column according to the manufacturer’s instructions (Waters). The eluted peptides were dried using a speed vacuum. Dried peptides were resuspended in 50% acetonitrile with 0.2% formic acid, and phosphopeptides were enriched using Fe-NTA resin. Peptides were incubated on Fe-NTA for 3-4 h and washed twice with 50% acetonitrile containing 0.2% formic acid. Finally, phosphopeptides were eluted using a 2.5% ammonium hydroxide buffer. An equal amount of phosphopeptides was used in LC/MS/MS analysis using the Thermo Orbitrap Fusion system. Fe-NTA resin was prepared by chelating Ni-NTA resin according to the method described by Sanford & Smolka, 2021, with slight modifications. After the mass spec run, raw data of phosphopeptides were analysed by Proteome Discoverer software from Thermo Fisher Scientific using two algorithms, Sequest HT and MS Amanda 2.0, with standardised consensus and processing workflow parameters with slight modifications, including an s/n ratio of 5, a threshold of 50, unique peptides, and static carbamidomethylation modification (+57.021 Da). Phosphosite detection was performed by selecting dynamic phosphorylation modifications for serine, threonine, and tyrosine. In general setting, precursor was selected as mass spectrometry 1(MS1) precursor and in spectrum properties filter, min precursor mass was 350Da, max precursor mass was 10,000 Da and minimum peak count was 1. In scan event filters, the mass analyser was Ion Trap Mass Spectrometry (ITMS), Fourier Transform Mass Spectrometry (FTMS), MS order was MSI, MS2, activation type was Higher-energy collisional dissociation (HCD), and Max collision energy was 1000. The maximum delta change for input was 0.05; target FDR (strict) was 0.01; target FDR (relaxed) was 0.05; and validation was based on q-value. We used the UniProt mouse database (uniport-mouse_2023_reviewed.fasta) with a maximum of 2 missed cleavages; the minimum peptide length was 6, and the maximum length was 144. We then calculated log2 fold change from abundance values detected by the software, and the calculated p-values using the one-way Chi-square test. We plotted the heat map for significantly differentially regulated genes using the SR plot web server, as described by Tang et al., 2023.

#### Statistical analysis

p-values were calculated using a two-sample t-test from the mean and standard deviation, as described by Xu et al. (2017), using the MedCalc online statistical software calculator (MedCalc Software, 2024).

## Author contributions

This work was completed in the lab of TM. TM initiated the project. TM and SS have designed the experiments and performed the analysis. DP has given suggestions on p-body markers. SS performed all the experiments and wrote the manuscript with input from all authors.

## Acknowledgements

The work was supported by the inStem core funds received by TM

## Materials availability

This study did not generate new unique reagents.

## Data availability statement

Phosphoproteomics data is submitted to the ProteomeXchange consortium MassIVE. The accession number for the dataset is MSV000099451. https://doi.org/doi:10.25345/C5NZ81358. No other dataset was generated in this study apart from phosphoproteomic dataset.

## Funding declaration

This work has not received any specific funding.

## Declaration of interests

The authors declare no competing interests

## Notes

### Competing Interest Statement

The authors have declared no competing interest.

### Summary of Updates

Updated gene names according to the nomenclature and added a few references.

