## Supplementary File S1_Supp Figures for "Calcium: Modulator of Post-transcriptional and post-translational process in mESCs"

**Supplementary Figures**

**Supp Figure S1**

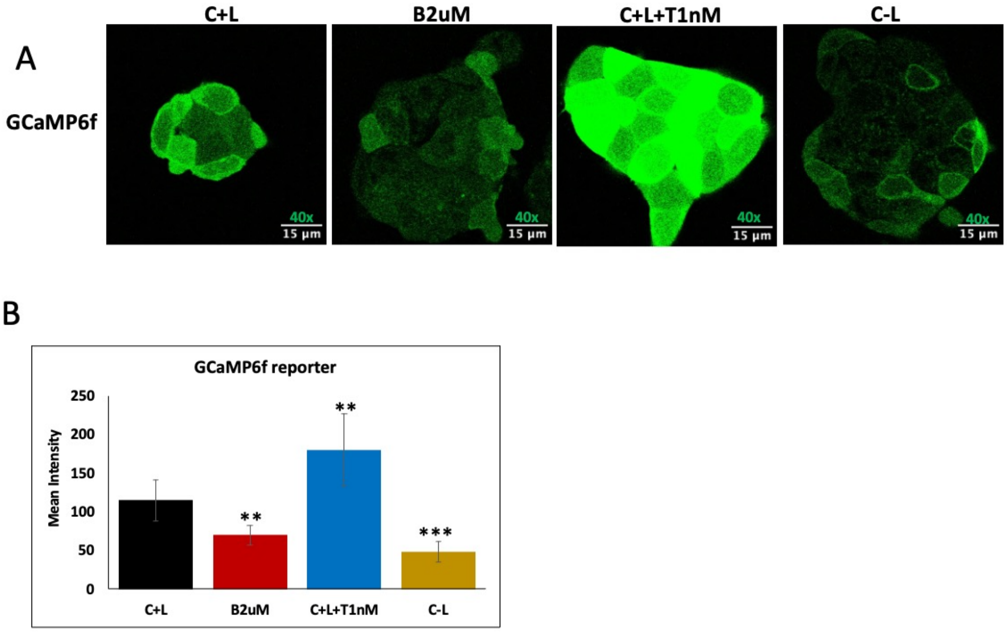


**Supp Figure 1: A) :** GCaMP6f reporter was used to check the intracellular $Ca^{2+}$ level in mESCs at different conditions (C+L, B2uM, C-L, C-L+T1nM) using excitation laser 488nm. (n=3) shown by representative images.

B) Bar plot represents the mean intensity of GCaMp6f reporter with B2µM, C+L+T1nM, and C-L compared to control C+L. The mean intensity was calculated from mESCs colonies containing approximately 11-31 cells from the confocal images (n=11) from different region of the above experiments

*C+L= Control media containing Lif; B2μM = C+L + BAPTA 2μM; T1nM= C+L + thapsigargin 1nM; C-L= control media without Lif; C-L+T1nM = C-L + thapsigargin 1nM; n = biological replicates. BAPTA and thapsigargin were used as calcium-modulating reagents.*

*The mean intensity was calculated from the confocal images from different regions for all experiments using ImageJ software.* An unpaired two-sample t-test was used to calculate p-value using the GraphPad online statistical software calculator (https://www.graphpad.com) with the mean and standard deviation. ***p-value representation with asterisk <0.05 (*), <0.01 (**), <0.001 (***), <0.0001 (****).***

**Supp Figure S2**

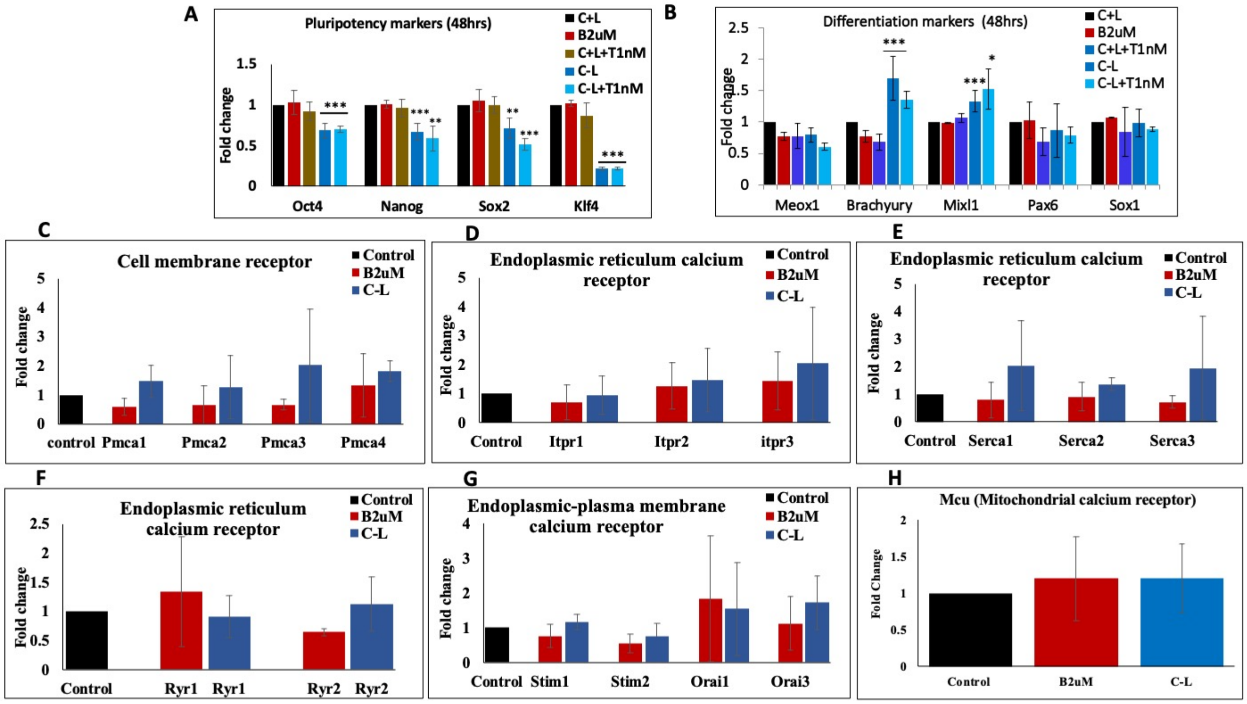


**Supp Figure S2: A, B)** QPCR of pluripotency (*Oct4, Nanog, Sox2 and Klf4)* and differentiation markers (*MeoX1, Brachyury, MixL1, Pax6 and Sox1*) of the mESCs cultured for 48hrs under different conditions (C+L, B2µM, T1nM, C-L, C-L+T1nM) (n=5).

**C, D, E, F, G, H)** QPCR of Plasma membrane calcium receptors (Pmca1, Pmca2, Pmca3, Pmca4), endoplasmic calcium receptor (Serca1,Serca2, Serca3, Ryanodine receptor 1 (Ryr1), Ryanodine receptor 2 (Ryr2), Itpr1, Itp2, and Itpr3), endoplasmic-plasma membrane receptors (Stim1, Stim2, Orai1 and Orai3) and mitochondrial calcium receptor (Mcu) in the mESCs cultured under different condition (control, B2µM, and C-L for 72hrs (n=4).

*C+L= Control media containing Lif; B2μM = C+L + BAPTA 2μM; T1nM= C+L + thapsigargin 1nM; C-L= control media without Lif; C-L+T1nM = C-L + thapsigargin 1nM; control= C+L+ DMSO equivalent to B2μM condition; n = biological replicates. BAPTA and thapsigargin were used as calcium-modulating reagents.*

An unpaired two-sample t-test was used to calculate p-value using MedCalc online statistical software calculator (MedCalc Software, 2024) with the mean and standard deviation. ***p-value representation with asterisk <0.05 (*), <0.01 (**), <0.001 (***), <0.0001 (****).***

**Supp Figure S3**

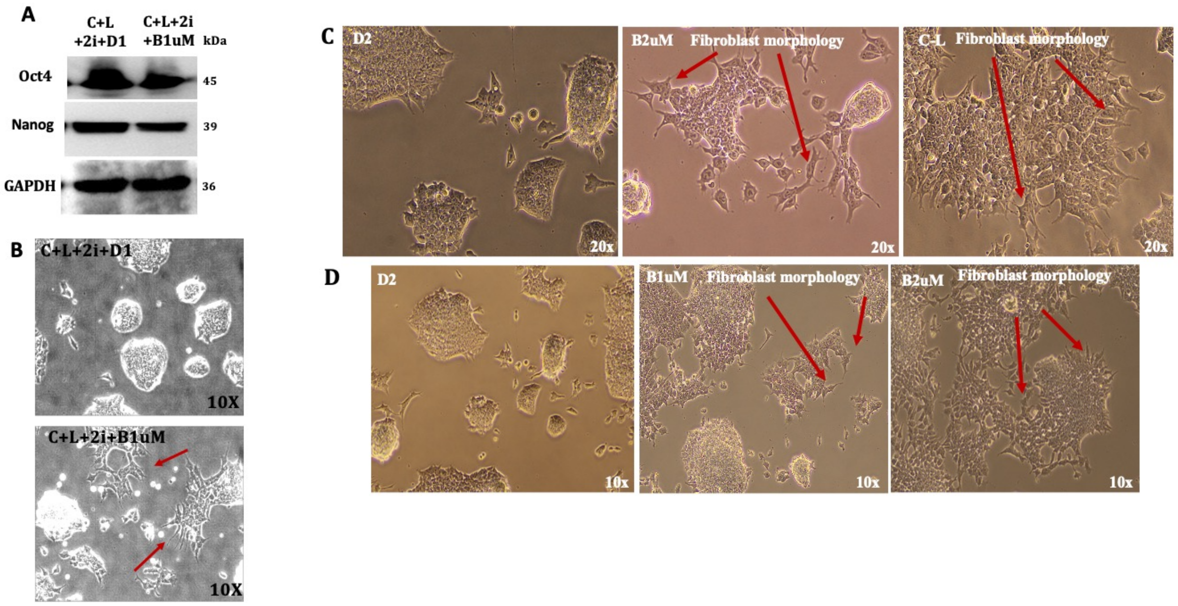


**Supp Figure S3: Legend: A)** Western blot detection of Oct4 and Nanog in the absence and presence of 2i inhibitor and B1uM (C+L+ 2i + B1µM). (n=3) shown by the representative images.

**B)** Brightfield images of the mESCs cultured in the absence and presence of 2i inhibitor and B1uM (C+L+ 2i + B1µM) at 20X magnification. (n=3) shown by the representative images.

**C, D)** Brightfield images of the mESCs cultured under different conditions (D2, B1um, B2µM and C-L) at 10x and 20x magnification. (n=4) shown by the representative images.

**Note:**

*C+L= Control media containing Lif; B1*μ*M= C+L + BAPTA 1*μ*M; B2*μ*M = C+L + BAPTA 2*μ*M; C-L= control media without Lif*; *D1= dmso equivalent to B1*μ*M condition, D2= C+L+ DMSO equivalent to B2*μ*M condition; C+L+2i inhibhitor = C+L+* CHIR 99021 (CH) (3μM) and PD0325901 (PD) (1μM); *n = biological replicates. BAPTA was used as calcium-modulating reagents.*

**Supp Figure S4**

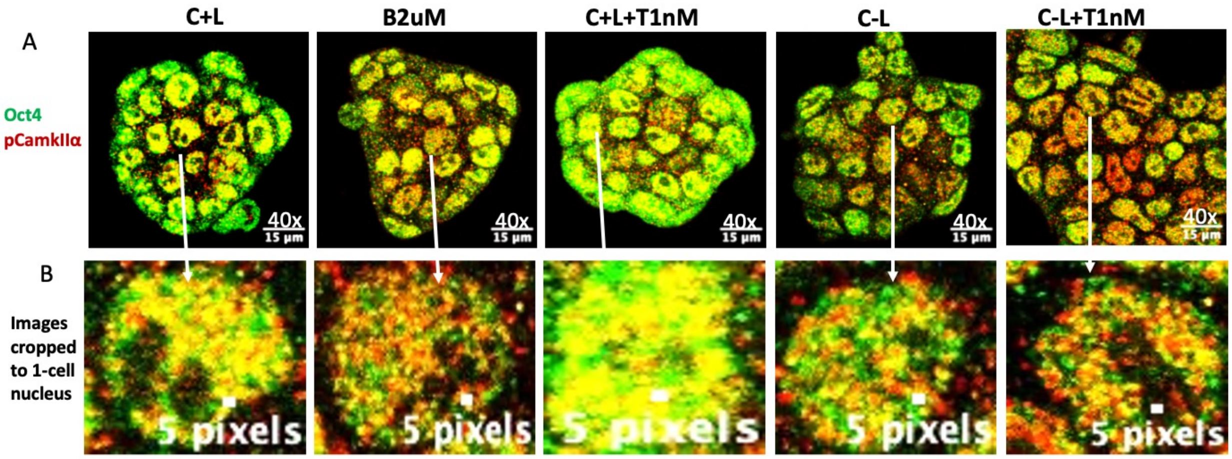


**Supp Figure 4 A)** Co-localization expression profile of Oct4 (green) and pCamkIIα (red) in mESCs cultured under different conditions (C+L, B2µM, T1nM, C-L, C-L+T1nM) (n=3) was observed from the composite images of Oct4 and pCamkIIα from Figure 2B, 2C. Co-localization of Oct4 and pCamkIIα is seen in yellow color.

B) The images obtained using a confocal microscope from the above experiment at 40x magnification were cropped to one nucleus size with a 5 pixels scale bar at the bottom to observe co-localization of Oct4 and pCamkIIα.

*C+L= Control media containing Lif; B1µM= C+L + BAPTA 1µM; B2µM = C+L + BAPTA 2µM; T1nM= C+L + thapsigargin 1nM; C-L= control media without Lif; C-L+T1nM = C-L + thapsigargin 1nM; n = biological replicates. BAPTA and thapsigargin were used as calcium-modulating reagents.*

**Supp Figure S5**

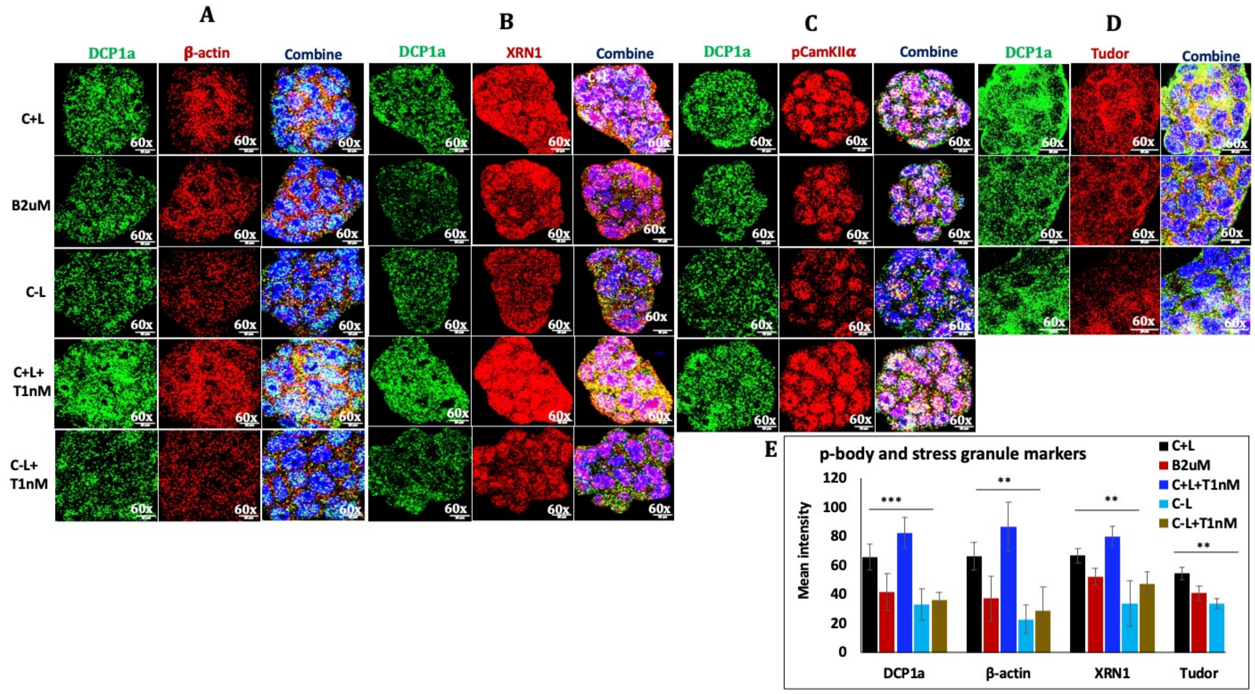


**Supp Figure S5: Legend: A)** IF detection of **A)** Dcp1a (green) and β-actin (red), **B)** Dcp1a (green), Xrn1 (red), **C)** Dcp1a (green) and pCamkIIα (red), **D)** Dcp1a (green) and Tudor (red) in mESCs cultured under different condition (C+L, B2μM, T1nM, C-L and C-L+T1nM) by confocal microscopy at 60X magnification. (n=3-4) shown by the representative images. IF experiments were done by thorough scanning the slide of different areas.

**E)** The bar graph represents the mean intensity of Dcp1a, Xrn1, pCamkIIα and Tudor expression in mESCs colonies containing approximately 11-20 cells, calculated from confocal images (n=15) from different regions from the above experiments (A).

*C+L= Control media containing Lif; B1μM= C+L + BAPTA 1uM; B2μM = C+L + BAPTA 2μM; T1nM= C+L + thapsigargin 1nM; C-L= control media without Lif; C-L+T1nM = C-L + thapsigargin 1nM; n = biological replicates. BAPTA and thapsigargin were used as calcium modulating reagents.*

*The mean intensity was calculated from the confocal images from different regions for all experiments using ImageJ software.* An unpaired two-sample t-test was used to calculate p-value using MedCalc online statistical software calculator (MedCalc Software, 2024) with the mean and standard deviation. ***p-value representation with asterisk <0.05 (*), <0.01 (**), <0.001 (***), <0.0001 (****).***

**Supp Figure S6**

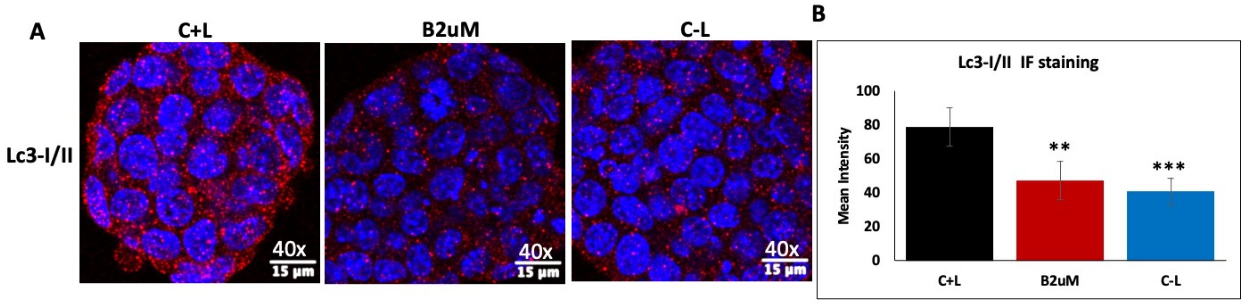


**Supp Figure 6** A) Immunofluorescence (IF) detection of Lc3-I/II in mESCs cultured under different conditions (C+L, B2μM, C-L) (n=3) using confocal microscopy at 40x magnification with 15um scale bar shown by the representative images. IF experiments were done by thorough scanning the slide of different areas.

B) The bar graph represents the mean intensity of LC3 expression in mESCs colonies containing approximately 20-27 cells, calculated from confocal images (n=11) from different regions of the above experiments (A).

*C+L = Control media containing Lif; B2μM = C+L + BAPTA 2μM; C-L= control media without Lif;* CaMKIIα-KD= CaMKIIα knockdown; *SER-R2-KD= SERCA-R2 receptor knockdown; n = biological replicates. BAPTA and thapsigargin were used as calcium-modulating reagents.*

*The mean intensity was calculated from the confocal images from different regions for all experiments using ImageJ software.* An unpaired two-sample t-test was used to calculate p-value using the GraphPad online statistical software calculator (https://www.graphpad.com) with the mean and standard deviation. ***p-value representation with asterisk <0.05 (*), <0.01 (**), <0.001 (***), <0.0001 (****).***

**Supp Figure S7**

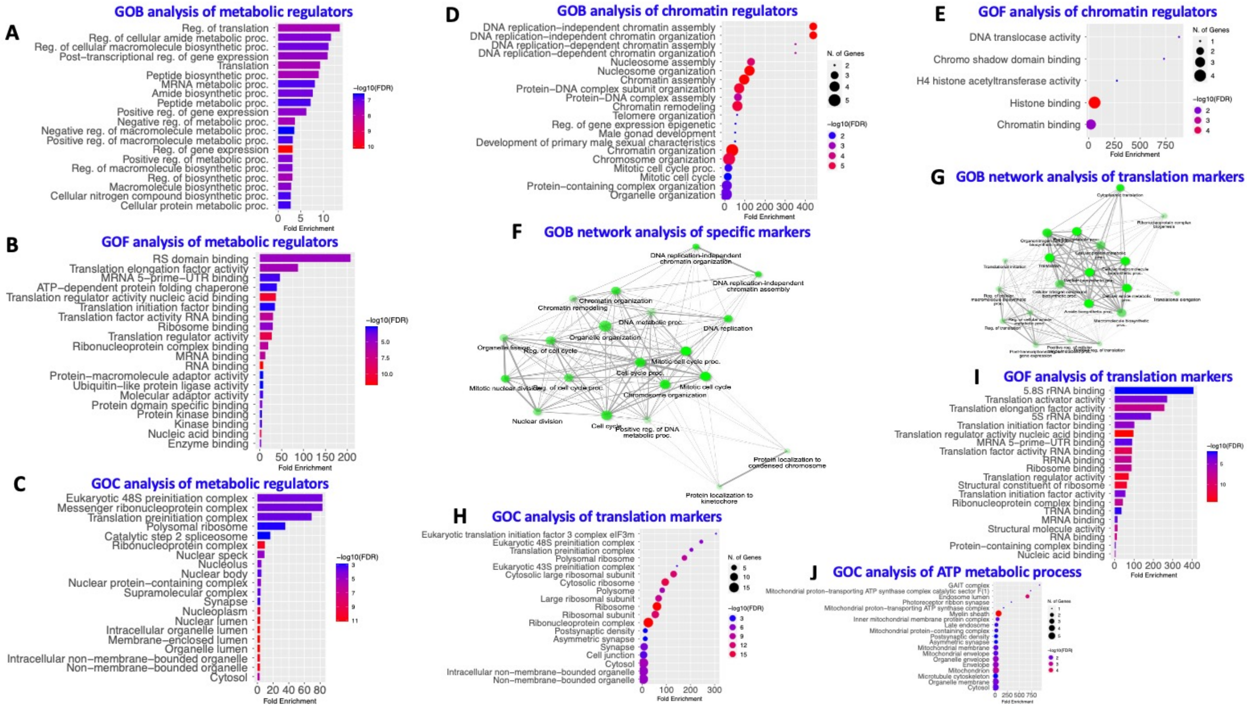


**Supp Figure S7: A)** GO biological process, **B)** molecular function, **C)** cellular component analysis was done for the DEPP involved in metabolic regulation including Hk2, Ranbp2, Trap1, Hspa8, Cdk1, Hint1, Snw1, Acin1, Gsk3b, Smarcd1, Pa2g4, Shmt2, Trip12, Abl2, Nmd3, Trim24, Eif3c, Gab1, Vac14, Nedd4, Akap12, Abcf1, Edc4, Stub1, Eef1a1, Hnrnpa1, Ddx21, Bclaf1, Eef2, Eif5a, Rbbp6, Nucks1, Eif2s1, Hsph1, Rpl5, Eef1d, Ssb, Eif3b, Ywhaq, Hnrnpab, Rbm39, Srsf5, Hnrnpa3, Hat1, Atrx, Gapdh, Nme1, Srsf6, Srsf1, Utf1, Tpx2, Tcof1, and Larp1 identified using string biological process analysis.

**D, E)** GOB, GOF analysis was done for the DEPP involved in chromatin remodelling identified using string biological process analysis.

**F)** GOB network analysis was done for DEPP including CSe1, smarcd1, Hk2, Pkm, Tfpt, Tcof1, champ1, nasp, Macf1, Hat1, Bcam, Atrx, Nmd3, Sde2, Ubr4, Hsph1, Trap1, nucks, Cdk1, Mki67, and Tpx2.

**G)** GOB network analysis was done for DEPP involved in translation regulation identified using string biological process analysis.

**H)** GOC, **I)** GOF analysis was done for the DEPP involved in translational regulation identified using string biological process analysis.

**J)** GOC analysis was done for DEPP involved in ATP metabolism identified using string biological process analysis.

**Supp Figure S8**

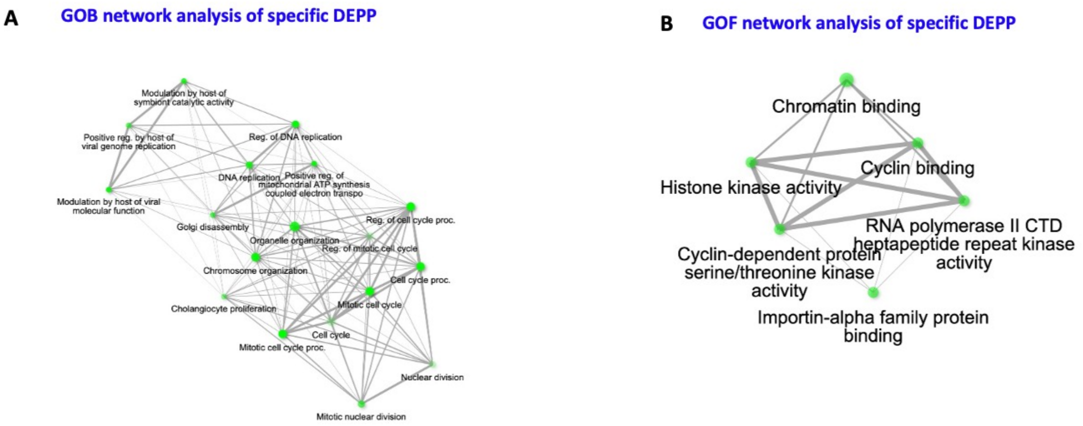


**Supp Figure S8: A)** GOB, **B)** GOF network analysis was done for DEPP including Cdk1, Mki67, Nucks1, and Tpx2.
